# Evolutionary dynamics of FoxQ2 transcription factors across metazoans: A tale of three ancient paralogs

**DOI:** 10.1101/2025.04.23.650082

**Authors:** Giacomo Gattoni, Che-Yi Lin, Joshua R York, Daniel Keitley, Carole LaBonne, Jr-Kai Yu, J Andrew Gillis, Elia Benito-Gutiérrez

## Abstract

FoxQ2 is a highly conserved class of Forkhead-box transcription factors expressed on the anterior side of the body in cnidarians and bilaterians. Despite this conserved expression pattern, recent phylogenetic analyses have revealed a complex and rapid evolution of this class, with several taxon-specific duplications and losses. Until recently, *FoxQ2* was thought to be lost in most vertebrate lineages, and its presence and localization in different vertebrate groups remains unclear.

To reconcile these conflicting reports of conservation and divergence, here we present a comprehensive analysis of the phylogenetic relationships and expression patterns of *FoxQ2* genes across metazoans. By combining phylogenetics and synteny analyses of *FoxQ2* sequences from 21 animal phyla, we uncover the presence of three ancient *FoxQ2* paralogs in bilaterians, which we name *FoxQ2a, FoxQ2b* and *FoxQ2c*. All three *FoxQ2* paralogs are present in the chordate lineage and two are conserved in vertebrates, indicating a richer repertoire of vertebrate Fox genes than previously estimated.

To investigate the expression of *FoxQ2* genes across bilaterians, we mined expression data from existing single cell transcriptomic datasets of mollusk, acoel, amphioxus and zebrafish development, and expanded it using fluorescent *in situ* hybridization in amphioxus, lamprey, skate, zebrafish and chicken. Our analysis demonstrates the conserved anterior expression of *FoxQ2a* and *FoxQ2b* paralogs while also revealing a novel domain of *FoxQ2c* expression within the chordate endoderm, including in amphioxus, lamprey and skate. Finally, we devise a method to predict conserved transcription factor binding sites across the three extant amphioxus genera with specificity to developmental stage and cell-type identity. This suggests conserved regulatory interactions for the expression of *FoxQ2a* across deuterostomes.

Overall, this work clarifies the complex evolutionary history of *FoxQ2* genes and identifies a newly discovered endodermally-expressed Fox gene, *FoxQ2c*. We further propose that the early duplication of *FoxQ2a* and *FoxQ2b*, along with their redundant functions, provided the ideal background for subfunctionalization, contributing to the fast evolutionary rate of *FoxQ2* sequences observed in bilaterians.

## Background

FoxQ2 is a member of the highly conserved Forkhead-box (FOX) family of transcription factors. The FOX family has ancient origins, likely tracing back to the common ancestor of Opisthokonta, and it has been identified in fungi, choanoflagellates and animals (1). Within the metazoan lineage, the repertoire of FOX proteins expanded dramatically to include ∼26 classes, named with letters from A to S (2). These proteins are characterized by the presence of a conserved forkhead or winged-helix DNA-binding domain and are involved in virtually all developmental processes as well as in metabolism and in the regulation of cell cycle (3,4). The forkhead motif is conserved in metazoans, composed of three alpha helices and three beta-sheets, but each class can be distinguished by subtle differences within and outside the DNA-binding domain (5–7).

The FoxQ2 class was initially described in the cephalochordate amphioxus, where phylogenetic analysis of a newly discovered *FoxQ* sequence revealed it belonged to a distinct group from *FoxQ1* genes (8). Since then, *FoxQ2* orthologs have been found in most animal phyla studied to date, and its evolutionary origin has been traced back to the ancestor of metazoans (9–11) (supplementary fig. 1). Recent studies have also highlighted the complex evolutionary history of this Fox class, marked by numerous taxon-specific duplications and losses, as well as rapid sequence divergence. Together, these factors complicate the understanding of how many paralogs are ancestral to specific groups (11–13). In contrast to this highly dynamic sequence evolution, the analysis of *FoxQ2* expression in a variety of invertebrates has revealed a remarkable conservation in the localization of *FoxQ2* transcripts within the anterior portion of the body (aboral for cnidarians) during early development (8,14–20). This expression pattern appears to reflect its function in the specification of anterior ectodermal identities, which has been investigated in detail only in a few deuterostomes (e.g. echinoderms) and protostomes (e.g. arthropods) (21–31). Notably, in many marine invertebrates FoxQ2 has been shown to be part of anterior gene regulatory network (aGRN) involved in the specification of the anterior neuroectoderm (16,28,32–37).

With the expansion of developmental analyses across an increasing number of taxa, the interest in the expression and function of *FoxQ2* has grown in recent years, though information about this conserved class remains fragmented. Here we comprehensively examine the complex evolutionary history and expression pattern of *FoxQ2* genes across metazoans, with a specific focus on chordates - the phylum in which *FoxQ2* was initially discovered but remains scarcely explored. Our phylogenetic analysis, which includes sequences from most animal phyla and is coupled with synteny analysis, identified three *FoxQ2* paralogs shared by bilaterians and cnidarians with numerous subsequent duplications and losses. We further characterize the expression of *FoxQ2* paralogs across bilaterians, focusing on different chordate lineages, revealing the existence of two ancestral *FoxQ2* genes in vertebrates. Finally, the here augmented information on *FoxQ2* expression and function is re-examined in the context of our new phylogenetic findings.

## Results

### High conservation but dynamic evolution of the *FoxQ2* class

To date, *FoxQ2* genes have been identified in species belonging to 14 phyla (supplementary fig. S1), including cnidarians (14,15,38), spiralians (mollusks, annelids, phoronids, brachiopods, nemerteans, orthonectida and platyhelminths) (16,17,19,20,39–42), ecdysozoans (arthropods, onychophorans and nematodes) (18,27,29,43) and deuterostomes (echinoderms, hemichordates, chordates) (8,13,24,32,44–47). Their presence in poriferans and ctenophores has also been suggested (48).

To expand the repertoire of known *FoxQ2* genes, and to further test the conservation of this class across Metazoa, we searched for orthologs in published genomes of taxa where *FoxQ2* had not been previously described. Among invertebrates, we found new *FoxQ2* orthologs in spiralians (gastropod mollusks, clitellate annelids, rotifers, bryozoans), ecdysozoans (tardigrada) and xenacoelomorphs. Additionally, we identified *FoxQ2* paralogs in most non-bilaterian metazoans, including poriferans, ctenophores and cnidarians (supplementary table S1). We then turned our attention to vertebrates, for which information on this gene was surprisingly scarce until recently, leading some researchers to speculate that this gene was lost in tetrapods or amniotes (11,12). Motivated by the recent discovery of *FoxQ2* orthologs in teleosts, coelacanths, sauropsids and monotreme mammals (49), we BLAST-searched the genomes of the lamprey *Petromyzon marinus* (50), the skate *Leucoraja erinacea* (51), and two amphibians, *Xenopus laevis* and *Pleurodeles waltl* (52). We found a single *FoxQ2* copy in the genomes of the lamprey and skate, while no ortholog was found in amphibians. Contrary to the previously accepted scenario, our search indicates that most vertebrate groups possess *FoxQ2* genes, and that these were secondarily lost in amphibians and placental mammals independently.

In sum, combining our BLAST-based search with sequences identified in previous publications, we retrieved candidate *FoxQ2* orthologs from 70 species belonging to 21 phyla. All of these contained forkhead domains clearly identified as belonging to the *FoxQ2* class (supplementary fig. S1A, supplementary table S1, see Materials and Methods). The number of *FoxQ2* paralogs in each species varies considerably both between and within phyla (supplementary fig. S1B). The genome of four sponge and two ctenophore species analyzed contained a single copy of *FoxQ2*, while most cnidarian species analyzed had more than one paralog. All xenacoelomorph species in our analysis had only a single copy. Within protostomes, all ecdysozoans analyzed only have a single *FoxQ2* copy, while in spiralians the number of paralogs is highly variable between and within each phylum, with the annelid *Paraescarpia echinospica* currently holding the record with 13 *FoxQ2* genes (11). Among deuterostomes, echinoderms have one or two copies and hemichordates vary from one to four, while most chordates have a single *FoxQ2* gene, with the exception of amphioxus that has three paralogs.

Despite the broad phylogenetic distances and the varying number of paralogs, we found the predicted secondary structures of FoxQ2 forkhead domains are highly conserved in deuterostomes, protostomes and xenacoelomorphs (supplementary fig. S2). In each species, the domain features a sequence of (fold – sheet – fold – fold – sheet – sheet – fold) with only minor variations in length and amino acid composition. This suggests that differences in the expression pattern and function across species and paralogs might be related to sequence changes outside the forkhead domain or to the regulation of *FoxQ2* gene expression.

### Phylogenetic and synteny analyses recover 3 *FoxQ2* paralogs in Bilateria

The highly variable number of *FoxQ2* paralogs in several phyla underscores the complex evolutionary history of the *FoxQ2* class. This raises the question of how many paralogs can be traced back to the common ancestors of metazoans, bilaterians, protostomes and deuterostomes respectively, and which paralogs instead evolved independently in specific lineages. To address these questions, previous studies have examined the phylogenetic relationships of Fox genes in cnidarians and bilaterians using an increasingly high number of species (11–13). Initially, the *FoxQ2* class was hypothesized to consist of two ancestral *FoxQ2* groups, distinguished by the presence and position of an Engrailed Homology 1 (EH)-i-like motif at the N-terminal or C-terminal of the protein (12,13). Accordingly, these groups were named *FoxQ2-N* and *FoxQ2-C*, or *FoxQ2* and *FoxQD* respectively. However, more recent analyses revealed that while *FoxQ2-C*/*FoxQD* proteins, with a C-terminal EH-i-like motif, appear to form a single monophyletic group, many other *FoxQ2* sequences are highly divergent, lacking an EH-i-like motif and distributing on multiple branches of the phylogenetic tree (11). This suggests a fast evolutionary rate, further evidenced by the presence of lineage-specific expansions that complicate the reconstruction of the evolutionary history of this class.

To reconcile these contrasting results, we aimed to reconstruct the phylogenetic relationship of *FoxQ2* genes with a more comprehensive taxonomic sampling, incorporating the new sequences we identified from bilaterians, cnidarians, ctenophores and poriferans. To this aim, we performed two parallel analyses. First, from our *FoxQ2* transcript database of 70 species, we selected those for which only complete *FoxQ2* sequences were available. This resulted in the analysis of sequences from 33 species belonging to 17 different phyla: 7 Spiralia, 3 Ecdysozoa, 3 Deuterostomia, Xenacoelomorpha, Cnidaria, Ctenophora and Porifera (fig. 1). To further expand the taxonomic sampling and include taxa for which only partial gene sequences were available, we also isolated and aligned the *FoxQ2* forkhead domain of 47 species from 21 animal phyla - 9 Spiralia, 5 Ecdysozoa, 3 Deuterostomia, Xenacoelomorpha, Cnidaria, Ctenophora and Porifera (supplementary fig. 3). Phylogenetic trees were constructed using Maximum Likelihood (ML) (fig. 1, supplementary fig. 3A) and Neighbour Joining (NJ) (supplementary figs. 3B-C) methods, using *FoxQ1*, *FoxL1* and *FoxP* as outgroups. All the sequences retrieved in this study branched within the *FoxQ2* clade, and the broad structure of the phylogenetic trees was highly robust across groups of sequences and phylogenetic methods, supporting their identification as *FoxQ2* orthologs (fig. 1, supplementary fig. 3). Moreover, the tree structure remained consistent when considering only deuterostomes or only protostomes (supplementary figs. 4A-B) and aligned with the unrooted tree and results from dimensionality reduction approaches (supplementary figs. 4C-D). Our novel and expanded analysis revealed the existence of three ancient groups of *FoxQ2* genes, all three present in bilaterians and cnidarians, that likely originated near the root of the metazoan tree. These three branches were recovered in all analyses, although their relative position on the tree varied depending on whether full sequences or forkhead domains were considered (supplementary fig. 3). We define these as three distinct *FoxQ2* paralogs named *FoxQ2a*, *FoxQ2b* and *FoxQ2c* (fig. 1).

**Fig. 1.**
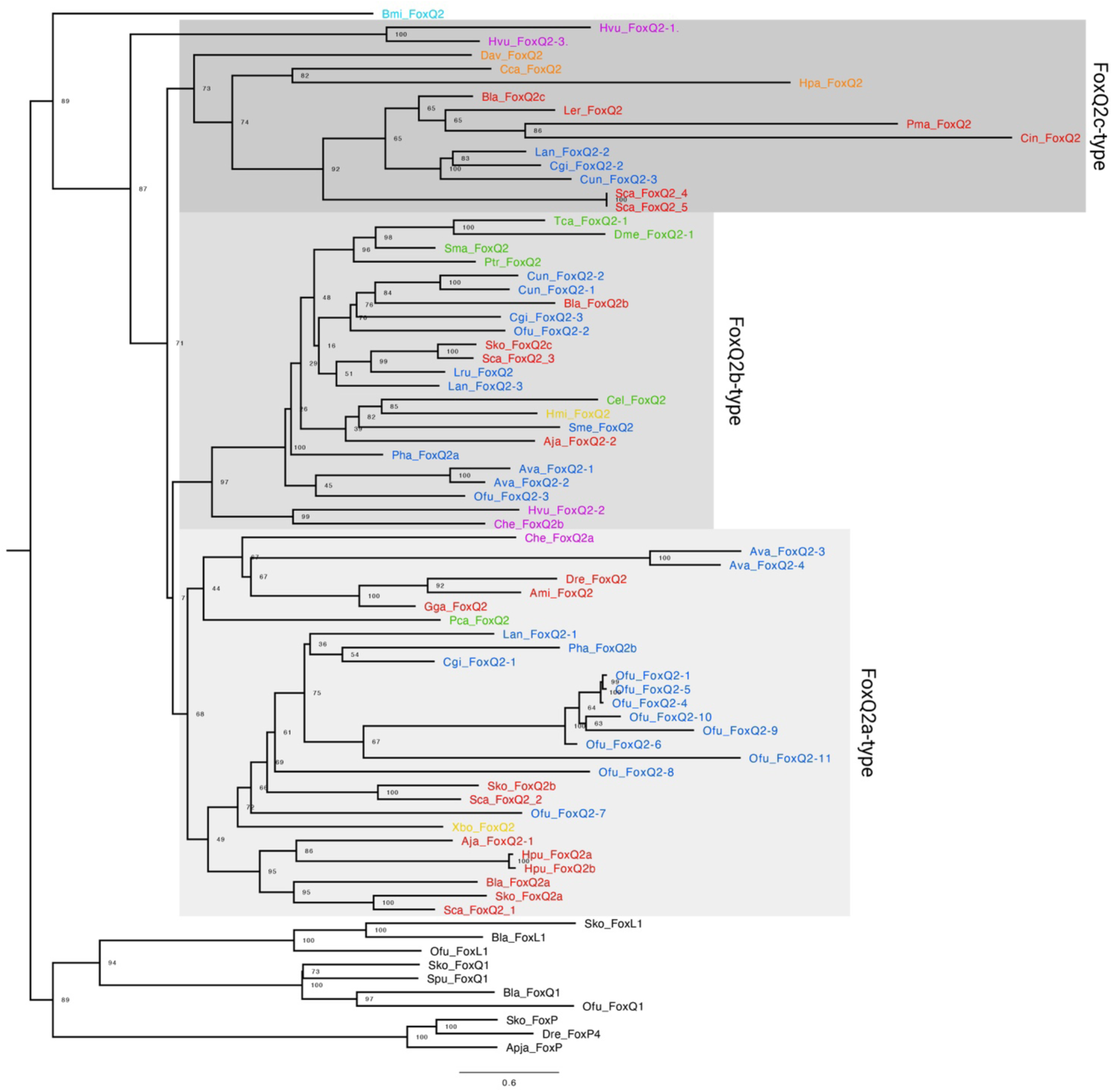
Phylogenetic analysis of metazoan FoxQ2 genes. Maximum likelihood tree topology based on *FoxQ2* complete gene sequences from 33 species belonging to 15 animal phyla. Individual genes are colored based on species taxonomy into deuterostomes (red), spiralian (blue) and ecdysozoan (green) protostomes, xenacoelomorphs (yellow), cnidarians (purple), poriferans (orange) or ctenoforans (cyan). Gray-shaded boxes demarcate the three *FoxQ2* types, *FoxQ2a*, *FoxQ2b* and *FoxQ2c*. *FoxL1*, *FoxQ1* and *FoxP* genes are used as outgroup.

The *FoxQ2a* family includes many fast-evolving genes previously identified as *FoxQ2-N* (*11*). Both ML and NJ trees retrieved *FoxQ2a* as a monophyletic clade, supporting previous analyses by Pascual-Herrera et al (12) (fig. 1, supplementary fig. 3). However, the main node had lower bootstrap values in most methods, suggesting fast sequence evolution and high divergence as proposed by Seudre et al (11). The family was identified in 12 metazoan phyla, including protostomes, deuterostomes, xenacoelomorphs, cnidarians and sponges (fig. 1, supplementary fig. 3A). *FoxQ2a* was present in most spiralians, including mollusks, annelids, brachiopods, phoronids and rotifers, often in high copy number, but was not recovered from flatworm, nemertean, orthonectida and bryozoan genomes. Moreover, its presence was variable even within spiralian phyla; for example, *FoxQ2a* was present in bivalve but not gastropod mollusks, and in several marine annelids but not in clitellates (supplementary table 1). Among ecdysozoans, *FoxQ2a* was found only in priapulids and was absent from all other lineages considered. However, the robust position of the priapulid sequence in all analyses supports *FoxQ2a* conservation in both protostome branches, followed by loss in most ecdysozoan lineages (fig. 1, supplementary fig. 3, 4). *FoxQ2a* orthologs were identified in all three deuterostome phyla, with independent duplications in hemichordates and sea urchin. Within chordates, the first *FoxQ2* gene discovered in amphioxus belonged to this clade, together with vertebrate *FoxQ2* orthologs in ray-finned fishes, reptiles and birds, while no paralog could be found in tunicates, cyclostomes and cartilaginous fishes (fig. 1).

*FoxQ2b* includes the highly-conserved *FoxQ2-C*/*FoxQD* genes that were recovered as a monophyletic clade in accordance with all previous analyses (11–13) (fig. 1). We identified *FoxQ2b* orthologs in 18 out of the 21 bilaterian phyla considered - with the exception of priapulids, ctenophores and poriferans - indicating a higher level of conservation of this gene compared to the other two *FoxQ2* families (supplementary table 1). Compared to *FoxQ2a*, *FoxQ2b* genes are generally present in lower copy number, with a single gene in each species and rare duplications in some annelid, mollusk and rotifer species. Despite the high conservation across bilaterians, it is also worth noting that within deuterostome phyla the gene has been lost in specific lineages, such as eleutherozoan echinoderms (starfish, sea urchins, sea cucumbers, brittlestars) and Olfactores (tunicates + vertebrates) chordates.

Surprisingly, our analysis identified a third, previously undescribed *FoxQ2* family: *FoxQ2c*. This family consistently branched separately from the other two in all phylogenetic analyses performed in this study (fig. 1, supplementary figs. 3, 4). When considering full sequences, both ML and NJ trees recover *FoxQ2c* as a sister clade to the rest of the class (*FoxQ2a* + *FoxQ2b*) with high bootstrap values (fig. 1). The forkhead domain analysis instead showed the *FoxQ2c* family nested within the tree as a sister clade to *FoxQ2b*, although with lower support (supplementary fig. 3A). While orthologs of the other two FoxQ2 families were found in most metazoan phyla analyzed, *FoxQ2c* was recovered only from specific lineages: brachiopods and mollusks among protostomes; chordates and hemichordates among deuterostomes; hydrozoan and schyphozoan cnidarians; and homoscleromorph sponges (fig. 1; supplementary fig. 3). Supporting this subdivision, a branch including *FoxQ2c*-type sequences from amphioxus and bivalve mollusks is also visible in a previous phylogeny of *FoxQ2* published by Seudre et al. (11). However, the low number of sequences in that study likely prevented its identification as a separate clade. Using additional sequences from cnidarians, mollusks, brachiopods, hemichordates, tunicates and vertebrates allowed this clade to be clearly visualized in our study. Among vertebrates, the FoxQ2c branch included only the cyclostome and cartilaginous fish sequences identified in this study, which branched separately from the *FoxQ2a* sequences found in bony fishes (fig. 1).

To confirm the presence of two separate *FoxQ2* paralogs in chordates, we performed synteny analysis of *FoxQ2* genes in four chordate species: amphioxus (*Branchiostoma lanceolatum*) (which possesses *FoxQ2* paralogs from all three families); zebrafish (*Danio rerio*) that has a *FoxQ2a*-type gene according to our analysis; lamprey (*Petromyzon marinus*) and skate (*Leucoraja erinacea*), which instead only have *FoxQ2c* genes (fig. 2A). Strikingly, we found that *FoxQ2c*-type paralogs are located in a similar genomic region in amphioxus, lamprey and skate, characterized by the close proximity to *FoxP* genes. Conversely, the conserved gene synteny of *FoxQ2a* previously identified in bony fishes (49) cannot be traced back to amphioxus (see below), further supporting the high divergence of this paralog.

**Fig. 2.**
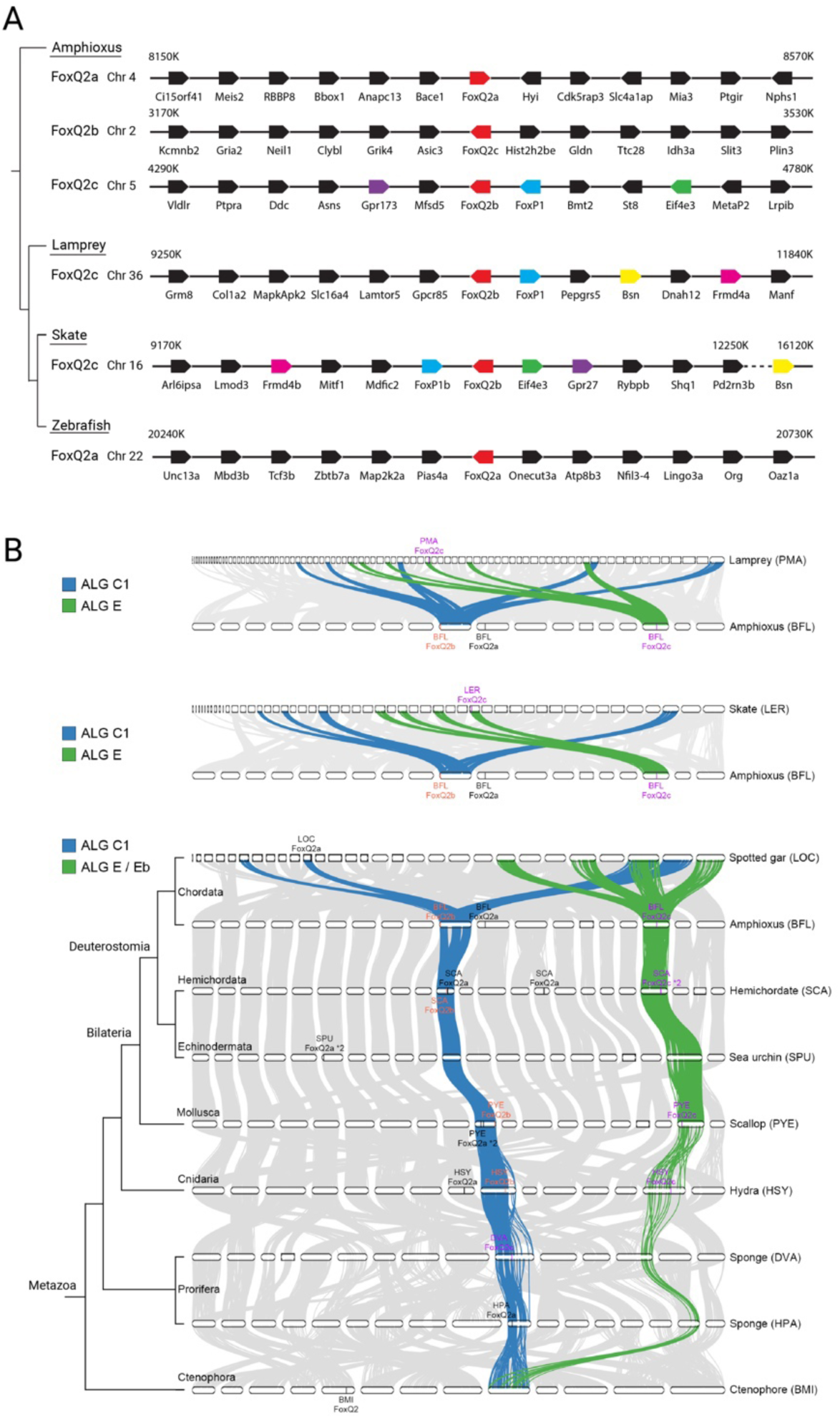
Synteny of FoxQ2 genes. **A.** Microsynteny analysis of the genomic environment around *FoxQ2* genes in chordates. Orthologous genes across species are indicated by color-coding the corresponding box. Chromosomal number and location of the area analyzed are indicated for each gene. **B.** Macrosyntenic orthology relationships of metazoan chromosomes in representative species of chordates, hemichordates, echinoderms, mollusks, cnidarians, ctenophora and porifera, highlighting two ancestral linkage groups where *FoxQ2a* and *FoxQ2b* (ALG C1) and *FoxQ2c* (ALG E/Eb) likely resided ancestrally.

In the three non-bilaterian phyla considered (Cnidaria, Ctenophora, Porifera) the assignment of *FoxQ2* sequences to *FoxQ2a-*, *FoxQ2b-* and *FoxQ2c*-types proved more challenging than in bilaterians, likely due to a high level of sequence divergence compared to bilaterian *FoxQ2* orthologs. Cnidarian sequences were found in all three *FoxQ2* branches, but in certain tree configurations *FoxQ2c*-types in *Hydra vulgaris* branched as sisters to all other *FoxQ2* groups (fig. 1, supplementary fig. 3). Similarly, in all analyses the ctenophore *FoxQ2* sequence branched separately at the base of the *FoxQ2* tree, indicating a particularly high sequence divergence from other *FoxQ2* sequences (fig. 1, supplementary fig. 3). For poriferans, the position of *FoxQ2* sequences varied based on the type of analysis: for example, *FoxQ2* from *Halichondria panicea* branched with *FoxQ2c*-type when considering full sequences (fig. 1), but branched with *FoxQ2a* when considering only the Forkhead domain (supplementary fig. 3A).

Recent comparative analyses of chromosome-level genomes from diverse bilaterians, cnidarians, sponges and ctenophores have revealed ancient chromosome-scale syntenies conserved across metazoan animals (53–55). We reasoned that these conserved macro-synteny patterns could provide valuable information for tracing the evolutionary history of *FoxQ2* genes, including the more elusive orthologs in basally-branching non-bilaterian metazoans, further strengthening and enhancing the results of our phylogenetic reconstruction. Given this, we performed macro-synteny analysis to compare the correspondence of ancestral linkage groups (ALGs) of *FoxQ2*-bearing chromosomes across 25 metazoan species spanning bilaterian, cnidarian, poriferan and ctenophore phyla (fig. 2B, supplementary table 2) (54). We found that *FoxQ2a*- and *FoxQ2b*-type genes were usually located in chromosomes originated from ALG_C1 (fig. 2B, chromosomes connected by blue ribbons), while *FoxQ2c*-type genes were located on chromosomes corresponding to ALG_E/Eb (fig. 2B, chromosomes connected by green ribbons) in chordates, hemichordates, mollusks, and cnidarians. Within the chordate lineage, vertebrates experienced dynamic gene loss of *FoxQ2* genes. For instance, we observed that the remaining *FoxQ2a* gene in spotted gar was still residing in ALG_C1-derived chromosome, while the surviving *FoxQ2c*-type genes in lamprey and skate are located on chromosome segments derived from ALG_E (fig. 2B, supplementary table 2). This is consistent with the notion that individual *FoxQ2* genes associated with different ALGs seem to have distinct evolutionary trajectories.

In poriferans, the *FoxQ2* sequences in the demosponges *Halichondria panicea* (HPA) and *Dysidea avara* (DAV) were found in ALG_C1-derived chromosomes (fig. 2B, supplementary table 2), supporting the hypothesis that ancestral FoxQ2 genes were already associated with ALG_C1 in the common ancestor of Porifera+Cnidaria+Bilateria. However, we found that *FoxQ2a-*type sequences in homoscleromorphs *Corticium candelabrum* (CCA) and *Oscarella lobularis* (OLO) were located on ALG_P-derived chromosomes (supplementary table 2), possibly due to a lineage-specific gene translocation event. In addition, we found that the ctenophore *FoxQ2* genes are located on a chromosome formed by the fusion of ALG_L and ALG_M. This genomic position was unique compared to those in all the metazoan genomes that we have examined, and thus could not allow us to distinguish between various possibilities regarding the position of *FoxQ2* genes in the metazoan common ancestor (see discussion). Therefore, although *FoxQ2* is present in this phylum, its relationship to the paralogs in other metazoans remains unclear.

The presence of *FoxQ2a* and *FoxQ2b* in the same ALG and the absence of a clear *FoxQ2b* paralog in early-branching metazoans (poriferans and ctenophores) also suggest that *FoxQ2a* and *FoxQ2b* share a closer evolutionary relationship, and they might have evolved by a tandem duplication event on ALG_C1 in the ancestor of Parahoxozoa (Cnidaria and Bilateria). The complementary distribution of *FoxQ2a* and *FoxQ2b* paralogs in ALG_C1-derived chromosomes in different species (supplementary table 2) may reflect an evolutionary process predicted by the duplication-degeneration-complementation model (56). Moreover, from this ancestral condition, *FoxQ2* paralogs have been considerably re-shuffled in selected lineages (supplementary table 2). Interestingly, we observed that *FoxQ2a*-type genes appeared to have undergone more frequent changes in their genomic position in various lineages, coinciding with their fast sequence evolution and high divergence in copy numbers among animals. For example, we identified duplicated *FoxQ2a*-type genes in both the hemichordate *Schizocardium californicum* (SCA) and the mollusk *Patinopecten yessoensis* (PYE) genomes. While in the scallop PYE, both *FoxQ2a*-type paralogs remained on the ALG_C1-derived chromosome, the *FoxQ2a*-type paralogs in the hemichordate SCA translocated to an ALG_O1-derived chromosome (fig. 2B, supplementary table 2). In echinoderms, *FoxQ2a* was translocated to ALG_H and ALG_O1 in sea urchin and sea star respectively, possibly indicating independent events in different echinoderm lineages. In amphioxus, *FoxQ2a* was translocated to a different ALG, ALG_A, which explains its different genomic location identified with microsynteny analysis.

Taken together, phylogenetic, microsynteny and macro-synteny analyses all support the presence of three ancient *FoxQ2* paralogs in metazoans, with two distinct groups (*FoxQ2a* + *FoxQ2b* and *FoxQ2c*) dating back to the ancestor of poriferans, cnidarians and bilaterians.

### Expression of *FoxQ2* paralogs across bilaterians: a conserved role in anterior development?

Following the first description of *FoxQ2* expression in the chordate amphioxus (8), its spatial distribution during development has been investigated in 12 phyla (supplementary fig. 5). The discovery of three *FoxQ2* paralogs allowed us to re-evaluate the expression pattern of *FoxQ2* genes previously reported in the literature for bilaterians. To the best of our knowledge, information on the localization of *FoxQ2a* is available for all deuterostome groups (chordates, echinoderms, hemichordates) (8,13,24,31,32,46,49,57) as well as annelids, mollusks and phoronids (11,19,39). *FoxQ2b* expression has been investigated in hemichordates, annelids, mollusks, nemerteans, brachiopods, platyhelminths, arthropods and onychophorans (11,13,16–18,20,28,29,39,42,43,58,59). Although these studies have rarely considered the phylogenetic relationships among *FoxQ2* genes, they generally suggest a conserved expression of *FoxQ2a-* and *FoxQ2b-*type genes in the anterior ectoderm. Conversely, the spatial expression of *FoxQ2c*-type genes has not been investigated in any species. By combining bulk RNA-seq, scRNAseq and *in situ* hybridization, here we provide new data on the expression of *FoxQ2a*, *FoxQ2b* and *FoxQ2c* paralogs across bilaterians.

As we hypothesized a high level of conservation of *FoxQ2a* and *FoxQ2b* anterior expression in bilaterians, we first queried publicly available scRNAseq datasets to detect the localization of *FoxQ2a* and *FoxQ2b* paralogs two species for which expression data is not available: the zebra mussel *Dreissena polymorpha* (bivalve mollusk) (60) and the acoel *Hofstenia miamia* (xenacoelomorph) (61) (supplementary fig. 6). *D. polymorpha* has 6 *FoxQ2* paralogs, including 4 *FoxQ2a*, 1 *FoxQ2b* and 1 *FoxQ2c*. We re-analyzed the data available for trochophore larvae, using the marker genes provided in the original publication to re-annotate the dataset (60), then plotted the expression of all six paralogs (supplementary fig. 6A). Of these, only one *FoxQ2a* paralog was expressed at significant levels, and we found that it labelled neuronal cells. The other *FoxQ2a* paralogs and *FoxQ2b* were instead detected only in scattered cells of the neural and ciliary ectoderm clusters. *FoxQ2* has not been investigated in the phylum Xenacoelomorpha, but the (debated) phylogenetic position of this group as the sister group to the other bilaterians makes it interesting to evaluate the evolutionary history of this gene (62). We found that the genome of *H. miamia* contains a single *FoxQ2* sequence belonging to the *FoxQ2b* clade (fig. 1). scRNAseq data of *H. miamia* hatchling juveniles is available as an interactive dataset. We therefore plotted the expression of *FoxQ2* (annotated as 98012160_foxb1 in the genome) and found expression in scattered cells within neoblast and neural clusters (supplementary fig. 6B). By referring to the spatial mapping described in the original paper (61,63), we observed that the neural clusters containing *FoxQ2*-positive cells were primarily located on the anterior portion of the acoel’s body.

We next turned our attention to chordates, where data on *FoxQ2* expression is scarce. In amphioxus, previous studies have shown that *FoxQ2a* starts to be expressed at the blastula stage, immediately following the maternal to zygotic transition, across the entire animal side of the embryo. Its expression then progressively restricts to the antero-dorsal side throughout gastrulation and neurulation (8,32,64). From late gastrula (G4) to early neurula (3-4ss) stages, *FoxQ2* is expressed in both neural and non-neural ectoderm, but by the mid neurula (7ss) stage its expression becomes restricted to the anterior epidermis, and at early larval stages it remains expressed at the tip of the rostrum and in a small portion of the mouth (32). Conversely, the expression of *FoxQ2b* and *FoxQ2c* has not been reported in the literature. By combining transcriptomic approaches and *in situ* hybridization chain reaction (HCR), here we show that in the European amphioxus *B. lanceolatum*, *FoxQ2b* starts to be expressed during the early phases of neurulation within the domain of *FoxQ2a* (fig. 3Ai, supplementary fig. 7A). Specifically, in the early neurula *FoxQ2b* is expressed in the most anterior portion of the neural plate and at the border between neural and non-neural ectoderm. As *FoxQ2a* restricts outside of the neural plate at the 7ss stage, *FoxQ2b* is also found only in the anterior epidermis, and in the larva it is co-expressed with *FoxQ2a* in the anterior rostrum (fig. 3Aii-iii). The expression of both *FoxQ2a* and *FoxQ2b* decreases during late development, as shown by RNAseq analysis (65) (supplementary fig. 7A,D). This data is corroborated by our analysis of a published developmental scRNAseq dataset of a different amphioxus species, *B. floridae* (66), which show early and widespread expression of *FoxQ2a* in neural and non-neural ectoderm, and a much more sparse and restricted expression of *FoxQ2b* (supplementary fig. 7B).

**Fig. 3.**
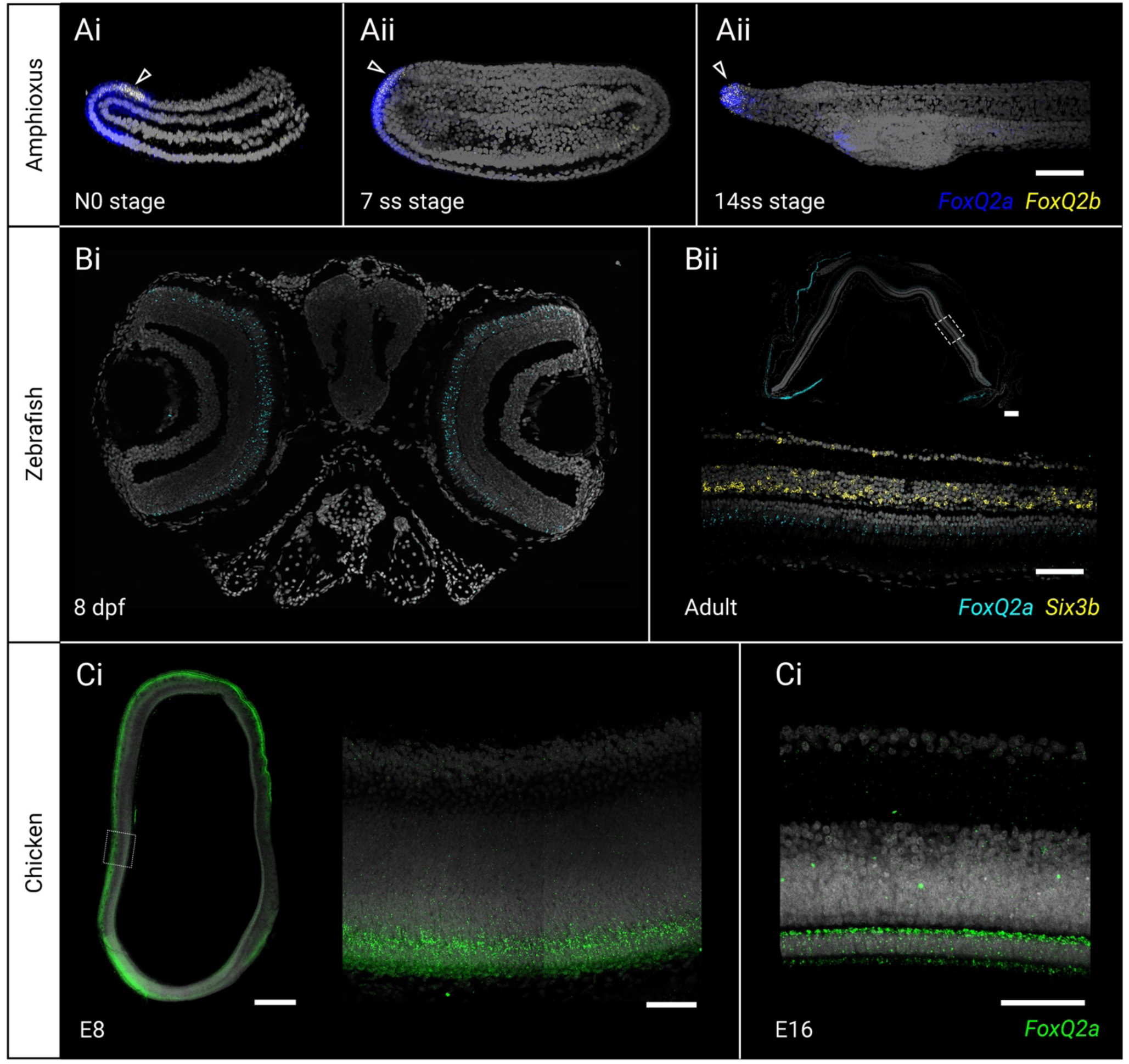
Expression pattern of *FoxQ2a-* and *FoxQ2b*-type paralogs in chordates by *in situ* hybridization chain reaction. **A.** Co-localization of *FoxQ2a* (blue) and *FoxQ2b* (yellow) in the anterior ectoderm of amphioxus whole-mount embryos spanning the key stages of neural tube development: N0 – early neurula (Ai); 7ss – mid neurula (Aii); 14ss – early larva (Aiii). Arrowhead indicates the area of co-expression of the two *FoxQ2* paralogs. **B.** Expression of *FoxQ2a* (cyan) in the retina photoreceptors of 8-day larvae (Bi) and adult (Bii) zebrafish, detected on paraffin sections of the larval head and adult eye respectively. Dashed square in Bii highlights the magnified section of the adult retina showing differential expression of *FoxQ2a* in the photoreceptor layer and *Six3b* (yellow) in nuclear and ganglionic layers. **C.** Distribution of *FoxQ2a* transcripts (green) in vibratome sections of the developing chicken eye at early (E8 – embryonic day 8, Ci) and late (E16 – embryonic day 16, Cii) stages, showing expression in the developing and mature photoreceptor layers of the retina. Dotted box in Ci indicates the position of the magnified section of the retina at E8 stage. Scalebars are 50µm for Ai-iii, Bi, Cii-iii, 100µm for Bii, 500µm for Ci.

Until recently, no information on the expression of *FoxQ2* was available for any vertebrate, and the gene was thought to be lost in several lineages. Although *FoxQ2a* orthologs had been detected in several species of ray-finned fishes, they were not found to be expressed during early zebrafish development, in contrast with its early distribution in many invertebrate deuterostomes. However, analysis of bulk RNA-seq and scRNAseq datasets that include late developmental stages shows that zebrafish *FoxQ2a*-type starts to be expressed after hatching in photoreceptor precursor and mature cells (supplementary fig. 7F) (67,68). A recent paper provided a detailed description of the expression and function of *FoxQ2* in zebrafish early larva (3-5 dpf) and showed that it is localized in blue cone cells, where it is essential to establish their identity (49). To date, this remains the only description of *FoxQ2a* expression in a vertebrate. By using *in situ* HCR, here we detected and compared the expression of *FoxQ2a*-type in later larval (8 dpf) and adult zebrafish as well as in chicken embryos (E8 and E16 stages) (fig. 3B-C). The analysis of zebrafish *FoxQ2a* confirmed its expression in the larval photoreceptor layer of the retina (fig. 3Bi), showing that it persists into adulthood and is thus not limited to development, while transcripts are absent in the brain (fig. 3Bii, supplementary fig. 8A). Turning to the chicken embryo, we found a very similar distribution of *FoxQ2a* in photoreceptor progenitors at E8 (fig. 3Ci), and later in the mature photoreceptor layer at E16 (fig. 3Cii), indicating a conservation in retinal expression of *FoxQ2a*-type genes across bony fishes.

### A novel domain of FoxQ2c expression in the endoderm

As indicated above, although at least four bilaterian phyla (mollusks, brachiopods, hemichordates, chordates) possess *FoxQ2c*-type genes, to the best of our knowledge no previous study has investigated their spatial expression. RNA-seq data across the development of bivalve mollusks and cephalochordates suggest that these genes are expressed between gastrula and larval phases (11,40,41,65): however, this does not provide information on the identity and location of *FoxQ2c*-positive cells. We therefore analyzed the localization of *FoxQ2c* in amphioxus (*B. lanceolatum*) using *in situ* HCR. Surprisingly, we found expression in a restricted domain within the endoderm of the early larva (12-14ss) which persisted at the one gill slit larval stage (fig. 4Ai-ii). In particular, *FoxQ2c*-positive cells are located within the midgut, just anterior to the intestinal thickening. This result is corroborated by recent scRNAseq data during development of *B. floridae*, showing *FoxQ2c* in the endoderm, and particularly in the midgut at the 14ss (T1) stage (supplementary fig. 7C) (66,69). While *FoxQ2a* and *FoxQ2b* expression decreases during late development, *FoxQ2c* remains active at high level in the endoderm even in the adult, where bulk RNA-seq shows it remains localized in gut tissues (supplementary fig. 7D) (65).

**Fig. 4.**
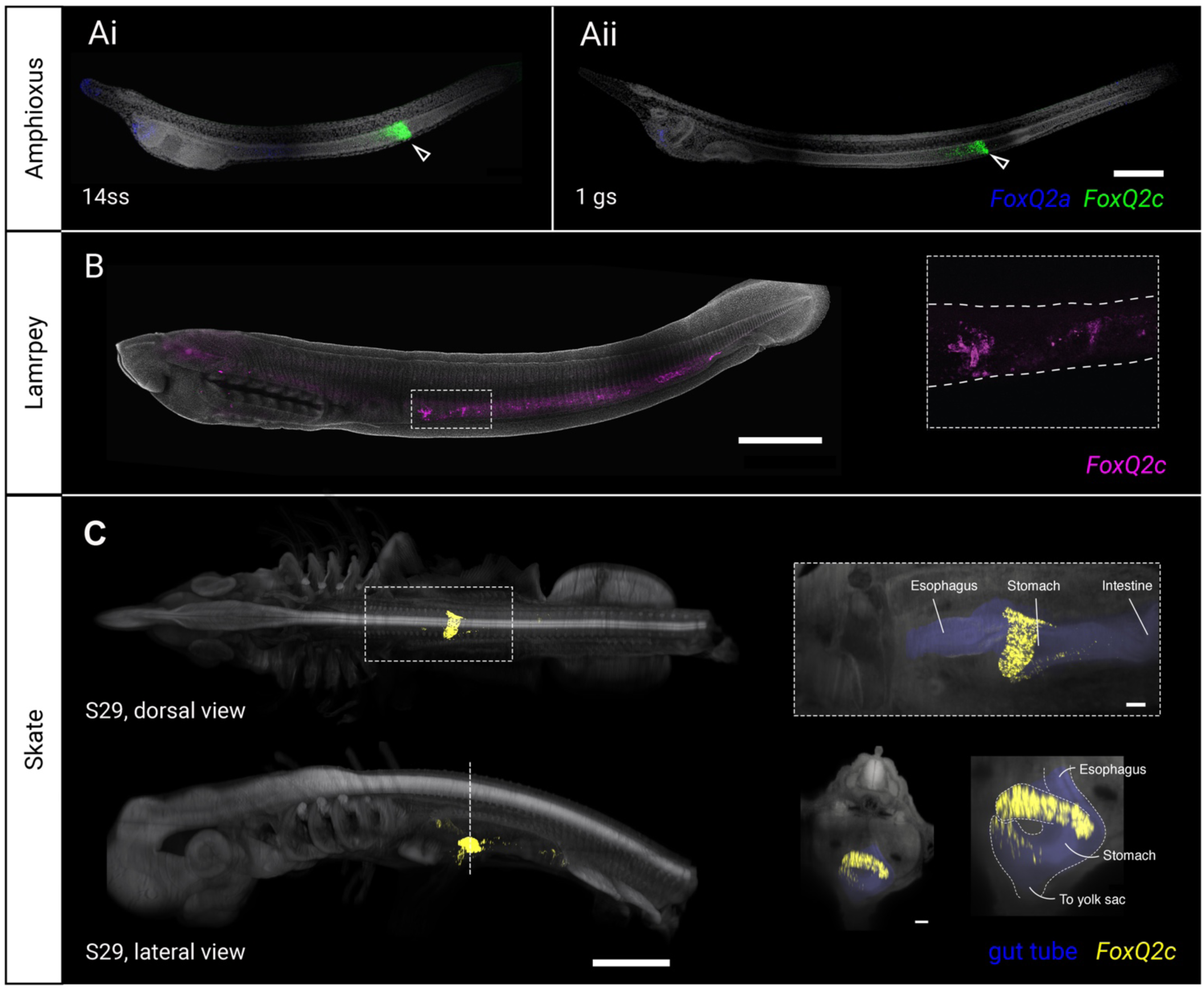
Expression of *FoxQ2c*-type genes in chordates. **A.** Co-detection of *FoxQ2a* (blue) and *FoxQ2c* (green) in amphioxus at early larva (14ss, Ai) and feeding 1 gill slit larva (1gs, Aii) stages. In the amphioxus larva *FoxQ2c* expression appears in a restricted portion of the midgut endoderm. **B.** Localization of *FoxQ2c* (magenta) in ammocoete larvae of the lamprey, showing expression in the gut (magnification in dashed box). **C**. Expression of *FoxQ2c* (yellow) in skate embyos at stage 29, showing restricted expression in a diverticulum of the digestive tube. Dashed box shows the magnification of the dorsal view, where the whole midgut is false-colored in blue. Dashed line indicates the level of the cross-section of the embryo, where the midgut is false-colored in blue. Scalebars are 100µm for A, 1mm for B, C.

Given the surprising localization of *FoxQ2c* in the amphioxus endoderm, we next sought to test whether this is an amphioxus-specific trait, or if *FoxQ2c* is also expressed in the endoderm of other chordates. To this aim, we investigated the expression of *FoxQ2c* in ammocoete larvae (40 days old) of the lamprey *P. marinus* and late embryos (S29) of the little skate *L. erinacea* (fig. 4B-C). *In situ* HCR on whole samples showed that *FoxQ2c* is indeed expressed in specific locations within the developing digestive system of basally-branching vertebrates: lamprey larvae had widespread expression across the middle and posterior portion of the gut tube, concentrated in scattered, strongly labelled cells (fig. 4B, supplementary fig. 8B). In skates, strong expression was found in the midgut, between the esophagus and the intestine, while more scattered *FoxQ2c*-positive cells are present at the base of the yolk stalk (fig. 4C, supplementary fig. 8C). Overall, these results strongly indicate a conserved midgut expression of *FoxQ2c* across chordates, defining a new domain of *FoxQ2* family expression.

To check whether *FoxQ2c* is expressed in the endoderm beyond chordates, we also looked at the expression of the *FoxQ2c*-type gene in the scRNAseq of *D. polymorpha* (60): at the stage analyzed we found only three positive cells, that were nonetheless located in the endoderm (supplementary fig. 6A). This could indicate that the gene is not expressed in the bivalve endoderm, or that it turns on at later developmental stages. In support of the second hypothesis, a previous RNA-seq analysis of adult *C. gigas* showed that *FoxQ2c*-type in this bivalve mollusk can be detected in the digestive gland (40), suggesting a conserved expression within endodermally-derived tissues. However, spatial analysis of expression in more mollusks and brachiopods is needed to test this hypothesis.

### Conserved candidate regulatory sequences across deuterostomes

Both *FoxQ2a* and *FoxQ2b* have been shown to be involved in the determination of anterior identity during embryonic development. Loss of *FoxQ2a* results in the loss of apical organ neurons in echinoderm larvae, of blue photoreceptors in zebrafish, and of anterior neural identity in the amphioxus cerebral vesicle, while loss of *FoxQ2b* in arthropod embryos causes defects in anterior brain development and labrum formation (22,25–27,29,31,69). Several studies have also shown that *FoxQ2a* and *FoxQ2b* genes are part of a highly conserved aGRN that is involved in the specification of anterior fate (70,32,35,16). However, little is known about the mechanisms that control *FoxQ2* expression. Functional studies and the analysis of cis-regulatory regions in echinoderms showed that *Meis* and *BicC* regulate *FoxQ2* maintenance, and *Six3/6* is required for *FoxQ2a* expression in sea urchin but not starfish (21,22,31,71–73). Similarly, *Six3/6* promotes *FoxQ2b* expression in arthropods (27,29).

Here, we aimed to compile a comparative list of candidate factors that regulate *FoxQ2* genes in cephalochordates, given that they are one of the few taxa to possess all three *FoxQ2* paralogs (fig. 5 shows the exemplified pipeline for *FoxQ2a*). To reconstruct conserved regulatory sequences across cephalochordates, we first identified *FoxQ2a*, *FoxQ2b* and *FoxQ2c* orthologs in five amphioxus genomes representing all three extant cephalochordate genera (*B. lanceolatum*, *B. floridae*, *B. belcheri*, *Asymmetron, Epigonichthys*). We then selected a region of ∼5000bp upstream of the start codon in all five species, and compared them for each gene using mVISTA (74) (fig. 5A). This approach identified conserved non-coding sequences (CNCSs) upstream of *FoxQ2* paralogs across cephalochordates: three CNCS for *FoxQ2a* (fig. 5A), one for *FoxQ2b* and one for *FoxQ2c* (supplementary fig. 9, supplementary table 3). In parallel, we used a published ATACseq datasets of *B. lanceolatum* at four stages of development, spanning early gastrula to larva stages (8hpf, 15hpf, 36hpf, 60hpf) (65), to identify open chromatin sequences. Strikingly, ATACseq peaks corresponded precisely to CNCS (supplementary fig. 9). These two lines of evidence supported their identification as conserved regulatory sequences. We therefore used CiiiDER (75) to predict transcription factor binding sites (TFBSs) within each CNCS for all three paralogs across species. To select only candidate conserved TFBSs, we normalized the length of each CNCS across species, and then selected only TFBSs that could be identified in all five species and located in similar positions (10% margin in each direction) (fig. 5B, supplementary table 3).

**Fig. 5.**
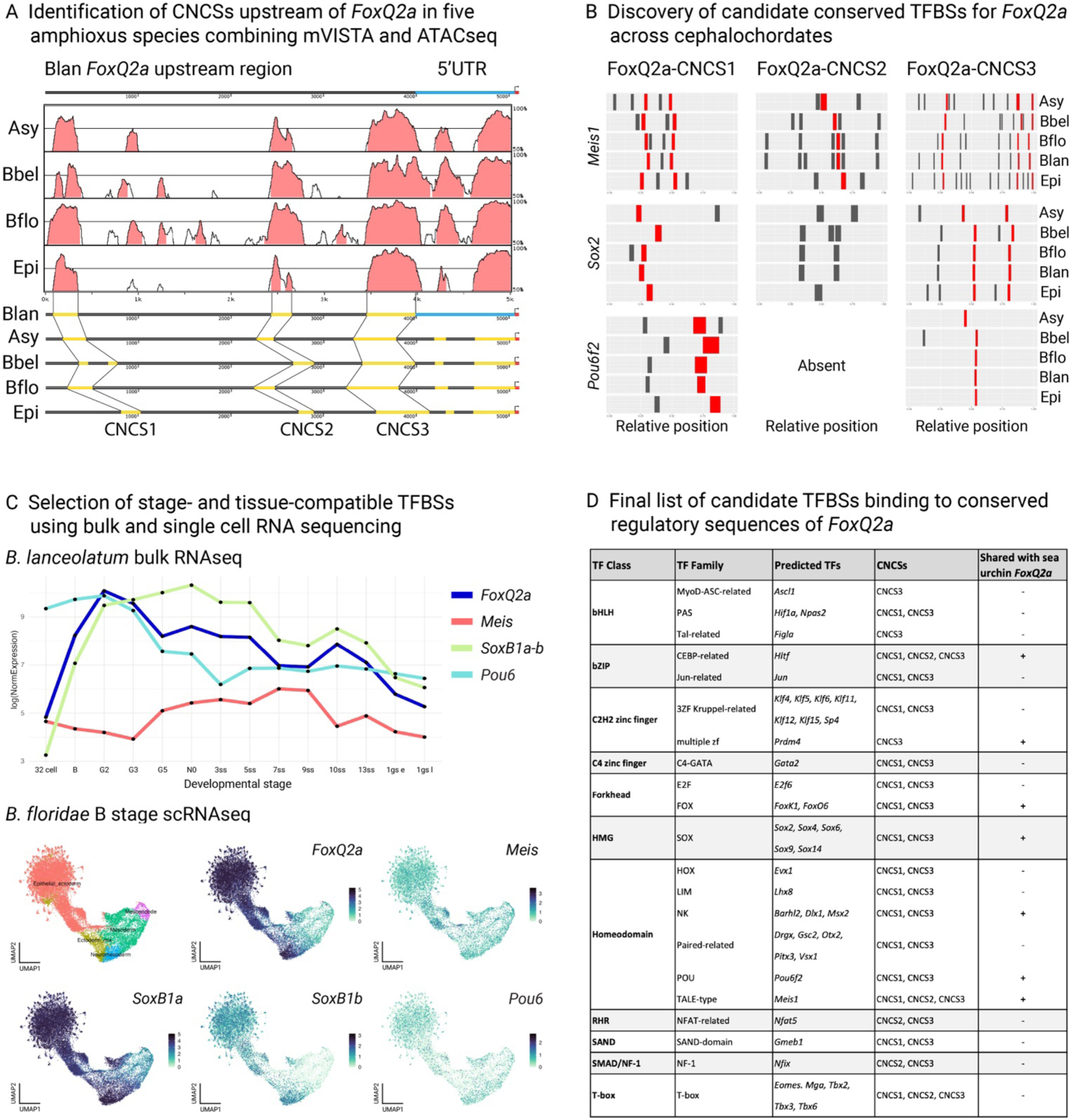
Prediction of conserved *FoxQ2* transcription factor binding sites with spatial and temporal resolution in amphioxus. Schematic workflow for the identification of transcription factor binding sites (TFBSs) for amphioxus *FoxQ2a*. **A.** Graphical representation of comparisons of the 5Kb upstream of *FoxQ2a* between *Branchiostoma lanceolatum* (Blan) and four other amphioxus species: *Asymmetron* (Asy); *Branchiostoma belcheri* (Bbel); *Branchiostoma floridae* (Bflo); *Epigonichthys* (Epi) using mVISTA, and schematic map of the position of the conserved non-coding sequences (CNCSs) in each species highlighted in yellow. **B.** Barplot showing the position of three predicted TFBSs (*Meis*, *SoxB* and *Pou6*) along the three CNCSs (x axis) found in the five amphioxus species (y axis). Sites that are present in all five species in the same position are marked in red. **C.** Developmental expression by bulk RNAseq allows to select among the list of conserved TFBSs those for transcription factors that are active when *FoxQ2* is expressed; further filtering with scRNAseq at the blastula stage, when *FoxQ2a* is first activated, retains only candidate TFBSs with the correct spatiotemporal resolution. **D.** Filtered list of conserved candidate *FoxQ2a* TFBSs in cephalochordates, highlighting the TF class and family, the presence in each CNCS and whether candidate TFBSs are shared with sea urchin *FoxQ2a* regulatory regions.

This analysis resulted in a list of candidate *FoxQ2* TFBSs in cephalochordates (supplementary table 3). We next refined this selection further by interrogating a published RNAseq datasets of *B. lanceolatum* development (65) and a recently published scRNAseq dataset of *B. floridae* development (69) (see Materials & Methods for details). This allowed us to select only candidate TFBSs for transcription factors that are active around the developmental time when each paralog is expressed, and in the correct cell type and embryonic location (fig. 5C, supplementary table 3). In *FoxQ2a* CNCSs we found, among others, multiple candidate TFBSs for transcription factors of the Meis, Pou6 and SoxB families, which were previously found in regulatory sequences of sea urchin and which are expressed maternally in both echinoids and amphioxus (71) (fig. 5B-D). As *Meis* is known to regulate *FoxQ2a* maintenance in sea urchin, these results suggest a possible conserved role of *Meis* in controlling *FoxQ2a* expression across deuterostomes. Putative TFBSs for *Meis* but not for *Pou6* and SoxB genes were found in the *FoxQ2b* CNCS, possibly suggesting regulatory differences that might underlie the difference in activation timing (supplementary table 3). Strikingly, the *FoxQ2c* CNCSs possessed TFBSs for endodermal markers, such as *FoxA* and *Pdx1*. Overall, this analysis provides a novel method for the identification of putative conserved and cell-type specific TFBSs, which can then be tested functionally, and suggests that *FoxQ2a* activation and maintenance in amphioxus might be under the control of early maternal signals that are conserved across deuterostomes.

## Discussion

### Evolution of the three *FoxQ2* genes across metazoans

*FoxQ2* is one of the most conserved classes of transcription factors in metazoans and has a widespread role in the specification of anterior ectodermal identity (28,32,33,35). At the same time, the *FoxQ2* family has undergone a dynamic evolutionary history, characterized by fast evolutionary rates and a high number of lineage-specific duplications and losses (11,12). Here, we specifically addressed this apparent discrepancy between conservation and divergence by analyzing the evolution of *FoxQ2* genes across all major metazoan lineages.

Our phylogenetic and synteny analyses, which included sequences from 21 animal phyla, revealed that the *FoxQ2* class originated in the common ancestor of metazoans, and identified three ancient *FoxQ2* paralogs, which we named *FoxQ2a*, *FoxQ2b* and *FoxQ2c*. Their evolutionary history is summarized in fig. 6. Through a survey of macro-synteny patterns across extant metazoan genomes, we found that *FoxQ2a*- and *FoxQ2b*-type genes are often located in chromosomes derived from ALG_C1, while *FoxQ2c*-type genes are mostly located on chromosomes corresponding to ALG_E/Eb. This trend is more apparent among bilaterian and cnidarian genomes, suggesting the existence of these two ancient FoxQ2 paralogs in the common ancestor of these animals. In addition, the presence of poriferan *FoxQ2a-*type and *FoxQ2c-*type genes in ALG_C1-derived chromosome also provides evidence to support the idea that in the common ancestor of poriferans, cnidarians and bilaterians, the original *FoxQ2* gene was likely located in ALG_C1. The initial gene duplication event generating the *FoxQ2c* paralog might have occurred in ALG_C1 as well. In contrast, the basal branching position of ctenophore *FoxQ2* sequence in the phylogenetic tree and its unique genomic position within the ALG_L/M group (fig. 1, fig. 2B) make it difficult to infer the ancestral position of *FoxQ2* gene in the common ancestor of all metazoans when we consider the ctenophore-sister hypothesis (55,76,77). If we consider the alternative sponge-sister hypothesis (78,79), the macro-synteny patterns would then suggest that the ancestral location of *FoxQ2* gene in the common ancestor of all metazoans was in ALG_C1, and that ctenophore *FoxQ2* was translocated to other ALGs and subsequently underwent drastic sequence divergence. Regardless these two competing phylogenetic hypotheses on whether sponges or ctenophores represent the sister group to all other animals, our results firmly support the existence of two distinctive groups of *FoxQ2* genes, namely the *FoxQ2a/FoxQ2b* and *FoxQ2c,* which can be traced back to the common ancestor of cnidarians and bilaterians. Following previous interpretations (11–13), we hypothesize that the ancestral *FoxQ2a/FoxQ2b* paralog further underwent additional duplication at the base of Parahoxozoa to give rise to *FoxQ2a* and *FoxQ2b*. These paralogs diverged extensively in the cnidarian and bilaterian lineages and their evolutionary relationships are therefore difficult to trace. Two less parsimonious alternative scenarios imply that only one *FoxQ2* gene was present in the ancestor of Parahoxozoa, and it independently duplicated in cnidarians and bilaterians, or that both paralogs were already present in the metazoan ancestor and *FoxQ2b* was lost in sponges and ctenophores. *FoxQ2a* and *FoxQ2b* paralogs were maintained in bilaterians, although subsequent independent events of gene loss resulted in different combinations of *FoxQ2* genes in each phylum (fig. 6). In spiralians, both paralogs were still present in most early lineages, but then underwent several taxon-specific expansions and reductions (11). In contrast, in ecdysozoans, *FoxQ2a* orthologs were lost in most lineages, leaving *FoxQ2b* as the only remaining member in arthropods, onychophorans, tardigrades, and nematodes. In deuterostomes, *FoxQ2a* was retained in all three lineages, while *FoxQ2b* was lost in eleutherozoan echinoderms and vertebrates, and both genes were lost in tunicates.

**Fig. 6.**
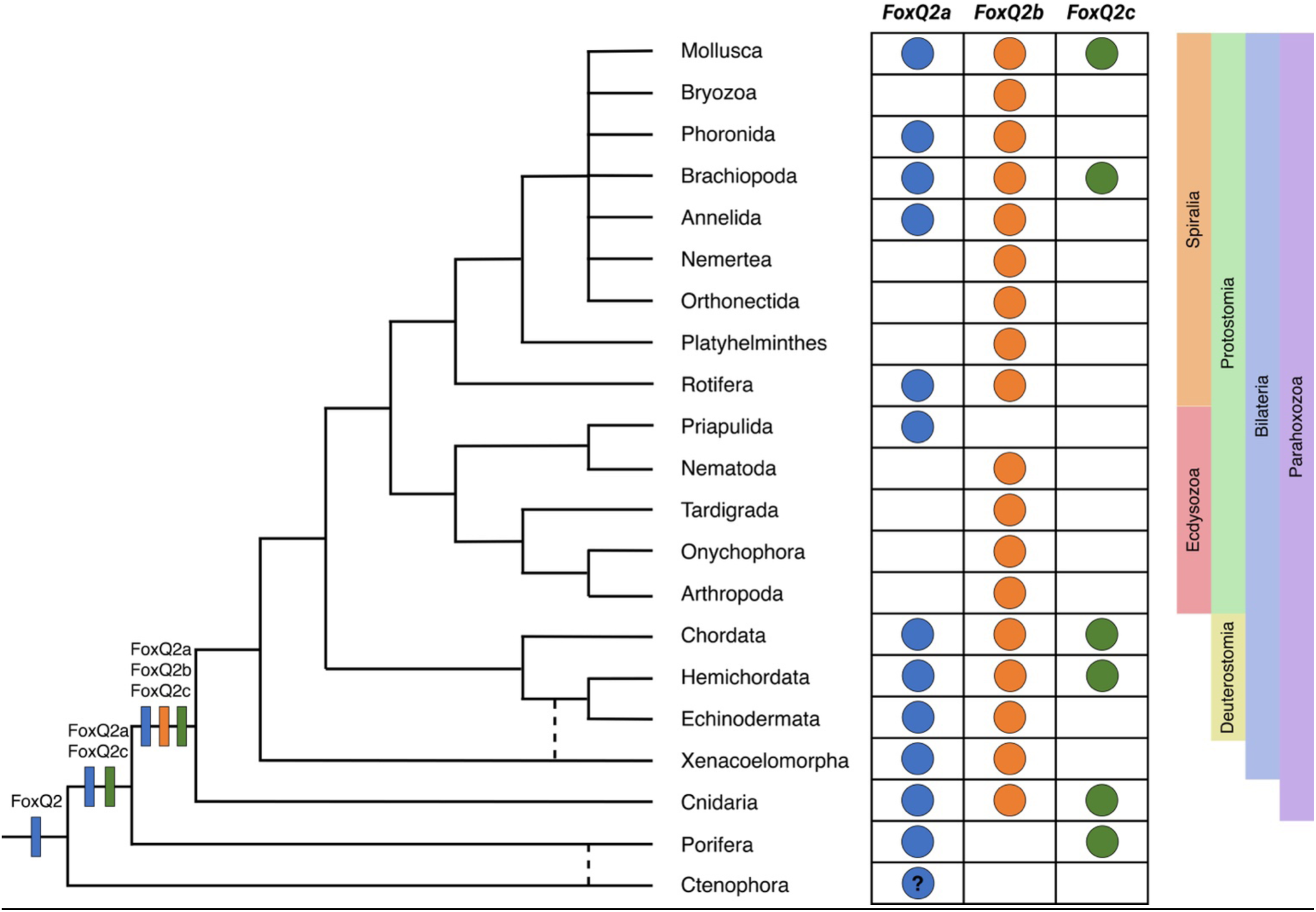
Schematic summary of the evolutionary history of *FoxQ2* genes in the main metazoan taxa.

The newly identified *FoxQ2c* paralog was lost in several lineages, including all ecdysozoans, the lineages leading to platyhelminths and rotifers within spiralians, and echinoderms among deuterostomes. However, it is still present in lophotrochozoans, hemichordates, and chordates. As such, cephalochordates, enteropneust hemichordates, bivalve mollusks and brachiopods are the only taxa in which all three paralogs have been identified in the same species to date. Focusing on chordates, the presence of *FoxQ2a*, *FoxQ2b* and *FoxQ2c* genes in amphioxus and of *FoxQ2a* and *FoxQ2c* genes in different vertebrate lineages, together with the conserved gene synteny of chordate *FoxQ2c*, indicates that vertebrates ancestrally possessed two *FoxQ2* genes. Cyclostomes and cartilaginous fishes then independently lost *FoxQ2a*, while bony fishes lost *FoxQ2c*. This, in turn, suggests that the vertebrate *FoxQ2* repertoire is richer than previously estimated and encourages further research on *FoxQ2* genes in this group.

The origin of both *FoxQ2a* and *FoxQ2c* at the base of metazoans, the ancestral presence of *FoxQ2c* on chromosomes corresponding to a separate ALG (ALG_E/Eb) compared to the other two paralogs, as well as the radically different expression pattern shown in this study, together call into question whether *FoxQ2c* should be considered a *FoxQ2* paralog or might belong to a new and separate family of forkhead transcription factors - with a highly similar Forkhead domain - that evolved independently from *FoxQ2a* and *FoxQ2b*. However, based on the data from ctenophores and combining all information from this phylogenetic analysis, we propose to rename *FoxQ2* paralogs into *FoxQ2a*, *FoxQ2b* and *FoxQ2c* (supplementary table 1). For example, as the amphioxus *FoxQ2b*- type has been previously annotated as *FoxQ2c* (and viceversa), we propose renaming both genes accordingly. For those species in which multiple copies of a paralog are present, the FoxQ2 type would be followed by a number. For example, *Saccoglossus kowalevskii* has two *FoxQ2a*-type copies and one *FoxQ2-b* copy originally called *FoxQ2-1*, *FoxQ2-2* and *FoxQ2-3* respectively. In the new nomenclature, these would be named *FoxQ2a-1*, *FoxQ2a-2* and *FoxQ2b*.

### Expression and function of *FoxQ2a* and *FoxQ2b* in bilaterians

In contrast with their ancient origin and high evolutionary divergence, *FoxQ2a* and *FoxQ2b* expression remained exceptionally conserved across Parahoxozoa. Orthologs of both genes are expressed and pattern the aboral ectoderm of cnidarians and the anterior ectoderm of bilaterians, with both neural and non-neural derivatives. Structures of the anterior neuroectoderm expressing *FoxQ2a* or *FoxQ2b* during development include: the apical organs present in the larvae of at least 8 phyla (14,16,17,19,39,46,47,57,80); the anterior portions of arthropod and onychoporan brains (18,27,28,43), planarian cerebral ganglia (42), amphioxus early neural plate (32), and hemichordate nerve plexi (13); and photoreceptor cells including the eyes of flatworms and chordates (39,58,81). Anterior non-neuroectodermal derivatives have also been described, such as the anterior epidermis and rostrum of amphioxus, the labrum of arthropods and the apical tuft cells of ciliated larvae (82,32,29,27). Interestingly, the expression dynamics of *FoxQ2a* and *FoxQ2b* genes in the anterior ectoderm appear to differ while remaining conserved across phyla. Indeed, *FoxQ2a* is generally expressed from very early stages, often at the beginning of zygotic transcription, in a broad animal/anterior domain, and then restricts towards the animal tip of the embryo. This pattern appears particularly conserved in deuterostomes, for which developmental data has been obtained for crinoid, echinoid and asteroid echinoderms, enteropneust hemichordates and cephalochordate chordates (23,24,31,32,46,57). After restriction, *FoxQ2a* remains expressed in many marine planktonic larvae and contributes to the specification of apical organ or apical tuft cells (22,23,26,57,70). Curiously, species that have lost their marine larva stage (vertebrate chordates, arthropods, panpulmonate mollusks, clitellate annelids) also appear to have lost either the early expression of *FoxQ2a* or the paralog altogether. There are however exceptions, such as polyclad flatworms, which have lost *FoxQ2a* but maintain a marine larva stage possessing an apical organ that expresses *FoxQ2b* (80). For species in which we have expression data of both *FoxQ2a* and *FoxQ2b*, such as annelids, hemichordates and cephalochordates, *FoxQ2b* appears after *FoxQ2a*, and generally starts to be expressed within the *FoxQ2a* domain (11,13).

The conservation in expression dynamics of *FoxQ2a* and *FoxQ2b* genes is reflected in their similar function within the aGRN that controls the specification of anterior neural identities. In fact, *FoxQ2a* mediates the formation of apical organ neurons and apical tuft cells in echinoderms, as well as the anterior brain in invertebrate chordates (22,32,81). Notably, a recent paper showed that *FoxQ2a* knock-out in amphioxus leads to the complete loss of anterior retina- and hypothalamus-like regions of the larval brain (cerebral vesicle) (69), supporting our previous hypothesis on the presence of a conserved anterior brain region, which forms in a Wnt-free area of the neuroectoderm through a conserved aGRN (32). Strikingly, loss of *FoxQ2a* in zebrafish leads to the disappearance of blue cones from the retina, which develops from the embryonic secondary prosencephalon (the portion of the central nervous system that includes telencephalon, hypothalamus and retina) (49). The similar expression of *FoxQ2a* in chicken embryos found here suggests that this is a conserved mechanism in multiple bony fish lineages. Another line of evidence comes from the fact that *FoxQ2a* was lost in placental mammals, concomitantly with the loss of blue photoreceptor types. However, the fact that amphibians, cyclostomes and cartilaginous fishes have lost *FoxQ2a* while still maintaining blue cones raises an intriguing hypothesis: that additional and partially redundant mechanisms may regulate blue cone formation in different vertebrate lineages.

Similar to *FoxQ2a*, *FoxQ2b* has been shown to control labrum and anterior brain formation (including central complex) in arthropods (27–29). Moreover, both paralogs appear to be negatively regulated by Wnt signalling, which originates from the vegetal pole and subsequently from the posterior side of the embryo during the development of most bilaterians. Wnt overactivation has been shown to downregulate expression of *FoxQ2a* in echinoderms, hemichordates and chordates, and of *FoxQ2b* in annelids and arthropods (16,23,28,32,70). This downregulation is followed by the loss of aGRN markers and by severe defects in the formation of anterior neuroectodermal structures, including larval apical organs in annelids and the anterior cerebral vesicle in cephalochordates. Strikingly, in cephalochordates, Wnt overactivation leads to a loss of both *FoxQ2a* and *FoxQ2b* (supplementary fig. 7E). These results suggest a similar and possibly redundant function for these two paralogs within the aGRN of bilaterians. However, it also indicates that functional considerations across bilaterians should only be made after careful examination, as the two genes have an ancient origin and have undergone independent evolution. This is further shown by the differences in the predicted TFBSs found in the proximal cis-regulatory regions of *FoxQ2a* and *FoxQ2b* across cephalochordates. By leveraging the availability of five amphioxus genomes from all three extant genera, as well as the genomics and transcriptomics resources build by the amphioxus community in the last decade, we devised a method for the predicting candidate conserved TFBSs with both developmental timing- and cell type-specificity. This resource reverals that while *FoxQ2a* regulatory regions contain candidate TFBSs that appear conserved with echinoderms (71), *FoxQ2b* is regulated by partially different transcription factors. Future analyses aimed at functionally testing the activity of these putative TFBSs could provide insights into the differences in the temporal activation of these two anterior *FoxQ2* paralogs in amphioxus.

### A newly-discovered endodermal *FoxQ2* gene

Finally, here we report the first expression pattern of the third *FoxQ2* paralog, *FoxQ2c*, in multiple chordate species. While the localization of the other two *FoxQ2* genes has always been associated with the anterior ectoderm, we find that this newly-described gene family is expressed in endodermal tissues during development. scRNAseq and *in situ* HCR in amphioxus, lamprey and skate show that *FoxQ2c* transcripts can be detected in the gut during late development. The timing of their activation suggests that this gene is not involved in early endodermal specification, but more likely in the differentiation of specific midgut cell types. In amphioxus, for example, *FoxQ2c* is activated at early larval stages, prior to the opening of the mouth and the start of feeding. These results suggest that chordates ancestrally possessed *FoxQ2c*-positive cell types in the midgut. Similar to the expression of *FoxQ2a* in the retina of only selected vertebrate lineages, the presence of *FoxQ2c* in the gut of amphioxus, lampreys and skates but not of other vertebrate lineages raises intriguing questions on the evolution of the endoderm. Were these cell-types lost in bony fishes? Or did their GRN modified extensively and lost *FoxQ2c* expression while maintaining the differentiated cell identity? The increasing availability of scRNAseq datasets from species across the chordate lineage will be critical to solve these questions and will help us gain a deeper understanding on the evolution of cell types.

## Conclusions

We have characterized the evolution of *FoxQ2* genes and identified three distinct paralogs. Two of these, *FoxQ2a* and *FoxQ2c*, are shared between poriferans, cnidarians and bilaterians, suggesting a more complex repertoire of forkhead genes in the metazoan ancestor than previously thought. Despite their high similarity in protein structure, these two ancient paralogs are expressed in distinct embryonic domains in bilaterians, i.e. the anterior ectoderm and the digestive endoderm respectively.

The third paralog *FoxQ2b* likely duplicated from *FoxQ2a* in the cnidarian-bilaterian ancestor, and the two genes are generally expressed in a similar domain in bilaterians. How does the fast molecular evolution of *FoxQ2a* and *FoxQ2b*, highlighted by phylogenetic analysis, reconcile with the high conservation in their expression pattern? We propose that the ancestral duplication of anterior genes *FoxQ2a* and *FoxQ2b*, which likely had a redundant function, might have provided an ideal background for subfunctionalization or specialization (65), so that new copies could be duplicated and others lost without large consequences for the organism. To test this hypothesis, it would be interesting to direct future studies at analyzing the function and possible redundancy of *FoxQ2a* and *FoxQ2b* paralogs in species that still possess both, such as annelids, mollusks, hemichordates or cephalochordates.

## Methods

### Phylogenetic analysis

Information on all sequences used in this study, including original gene name, new proposed name, species, taxon, paralog type, completeness, protein sequence, reference genome and reference code (when available), is provided in supplementary table 1. We first selected sequences from 17 phyla already available in the literature; incomplete sequences for *Gallus gallus FoxQ2* and *Branchiostoma lanceolatum FoxQ2c* were manually expanded from the genome. We then used reciprocal BLAST best hits to recover *FoxQ2* sequences from the genomes of *Petromyzon marinus, Leucoraja erinacea*, *Pleurodeles waltl* (Chordata), *Schizocardium californicum* (Hemichordata), *Candidula unifasciata, Mercenaria mercenaria* (Mollusca), *Lombricus rubellus*, *Hirudo medicinalis* (Annelida), *Membranipora membranacea* (Bryozoa), *Adineta vaga* (Rotifera), *Hypsibius exemplaris* (Tardigrada), *Priapulis caudatus* (Priapulida), *Hofstenia miamia*, *Symsagittifera roscoffensis*, *Xenoturbella bocki* (Xenacoelomorpha), *Hydractinia symbiolongicarpus*, *Hydra vulgaris*, *Acropora millepora*, *Pocillopora verrucosa*, *Rhopilema esculentum* (Cnidaria), *Dysidea avara*, *Ephydatia muelleri*, *Halichondria panicea*, *Corticium candelabrum*, *Oscarella lobularis* (Porifera), *Bolinopsis microptera*, *Hormiphora californensis* (Ctenophora). For all sequences used in the analysis, membership to FoxQ2 family was assessed by confirming the presence of a FoxQ2 domain using NCBI conserved domain search based on Conserved Domain Database (CDD) v3.21 (83). Following this confirmation, two groups of sequences were used for phylogenetic analysis:

- Group 1: complete *FoxQ2* sequences from 33 species belonging to 17 phyla,
- Group 2: sequences in which the *FoxQ2* domain was specifically isolated from 47 species belonging to 21 phyla.

For both groups, amino acid sequences were aligned using MAFFT v.7.526 under default settings (84). Phylogenetic analysis was then performed for both groups with two methods: Maximum Likelihood, using IQ-TREE web server with default parameters (85), and Neighbor Joining using Seaview (86). A thousand ultrafast bootstraps were used to extract branch support values with each method. The resulting trees were then visualized with FigTree v.1.4.4. UMAP dimensionality reduction and visualization was performed on the web version of Alignmentviewer (https://alignmentviewer.org/).

Microsynteny analysis was performed by manually comparing the genomic loci around *FoxQ2* genes in *Branchiostoma lanceolatum* (v.klBraLanc5.hap2, GCF_035083965.1), *Petromyzon marinus* (v.kPetMar1.pri, GCF_010993605.1) (50), *Leucoraja erinacea* (v.Leri_hhj_1, GCF_028641065.1) (51) and *Danio rerio* (v.GRCz11, GCF_000002035.6) in NCBI Datasets.

### Macro-synteny

Protein sequences and gene models were obtained from publicly available sources, with genome data details provided in supplementary table 2. Candidate FoxQ2 genes were identified using OrthoFinder v2.5.4 (87), and their classification was based on the results of phylogenetic analysis. Pairwise macrosyntenic comparisons between species were conducted using MCscan (Python version) implemented in JCVI v1.2.7 (88,89), as previously described (90,91). Briefly, Orthologous gene pairs between species were identified using the LAST aligner integrated in MCscan. A C-score threshold of 0.99 was applied to retrieve reciprocal best hits. For comparisons involving amphioxus and species that have undergone whole genome duplication (WGD), including lamprey, skate, and spotted gar, a relaxed C-score threshold of 0.7 was applied. Corresponding chromosome pairs were determined using Fisher’s exact test with Bonferroni correction (adjusted p-value < 0.05). To enhance the sensitivity in detecting syntenic relationships between sponge and ctenophore, a more lenient adjusted p-value threshold of 0.2 was used. Gene pairs located outside the identified corresponding chromosome pairs, as well as those involved in a small-scale chromosomal rearrangement event in hemichordate (91), were excluded from macro-synteny visualizations.

### Animal collection

#### Amphioxus

Adult *Branchiostoma lanceolatum* were collected in Banyuls-sur-Mer (France) and maintained in a custom-made facility at the Department of Zoology, University of Cambridge (UK). Spawning and fertilization was performed following (92), and embryos were raised in Petri dishes in filtered artificial salt water at 21°C. For embryos at 4 dpf, from the 48 hours stage embryos were fed with a mix of algae. At the desired stage embryos were collected and fixed in ice-cold 3.7% Paraformaldehyde (PFA) + 3-(Nmorpholino) propane sulfonic acid (MOPS) buffer for 12 hours, then washed in sodium phosphate buffer saline (NPBS), dehydrated and stored in 100% methanol at - 20°C.

#### Zebrafish

Zebrafish (*Danio rerio*) embryos were raised to 8 dpf and fixed overnight at 4°C in 4% paraformaldehyde in phosphate-buffered saline (PBS), then rinsed in PBS, dehydrated into 100% methanol and stored at -20°C. A Experiments using larval and adult zebrafish were conducted according to protocols approved by the Institutional Animal Care and Use Committees in facilities accredited by the Association for Assessment and Accreditation of Laboratory Animal Care International (AAALAC). The use of tissue samples for this study was reviewed and ethically approved by the University of Cambridge Animal Welfare and Ethical Review Body (AWERB) Committee.

#### Chicken

Fertilized White Leghorn Chicken (*Gallus gallus*) eggs (Charles River) were incubated in a 38°C humidified chamber with embryos staged according to (93). At E8 and E16 stages, eyes were dissected from the embryo in cold PBS and immediately fixed overnight at 4°C in 4% paraformaldehyde in PBS, then dehydrated to 100% methanol and stored at -20°C.

#### Lamprey

Adult sea lamprey (Petromyzon marinus) were collected from the Hammond Bay Biological Station, Millersburg, MI, and shipped to Northwestern University. Embryos and larvae were obtained by in vitro fertilization, fixed for 2h at room temperature in MEMFA at desired stages, and then dehydrated and stored in 100% methanol at -20°C prior to analysis. All procedures were approved by Northwestern University’s Institutional Animal Care and Use Committee (IACUC A3283-01).

#### Skate

Skate (L. erinacea) embryos were obtained from the Marine Resources Center at the Marine Biological Laboratory (MBL) in Woods Hole, MA, U.S.A., and reared to stage 29 as described in (94). All skate experiments were conducted according to protocols approved by the Institutional Animal Care and Use Committee of the MBL. Skate embryos were euthanized with an overdose of MS-222 (1 g/L in seawater), and all embryos were fixed overnight at 4°C in 4% paraformaldehyde in phosphate-buffered saline PBS, and then rinsed in PBS and dehydrated into methanol prior to analysis.

### In situ hybridization chain reaction

For *in situ* HCR v.3 (95) on amphioxus, zebrafish, chicken and skate specimens, probes were ordered through Molecular Instruments Inc.

#### Amphioxus

Reactions were performed as described in (96). Amphioxus embryos and larvae were rehydrated in NPBS + 0.1% Triton X, (NPBT), bleached for 30 minutes (5% Deionized formamide, 1.5% H2O2, 0.2% SSC in nuclease-free water) and permeabilized for 3 hours (NPBS + 1% DMSO + 1% TritonX). The embryos were incubated in Hybridization Buffer (Molecular Instruments) for 2 hours and then probes were added overnight at 37°C. The following day, embryos were rinsed in Wash Buffer (Molecular Instrument) followed by 5X-SSC + 0.1% Triton X. The samples were then incubated in Amplification Buffer (Molecular Instruments) for 30 minutes and left overnight in the dark at room temperature in Amplification Buffer + 0.03µM of each hairpin (Molecular Instruments). Embryos were washed in the dark in 5X-SSC + 0.1% Triton X and incubated overnight with 1 µg/mL DAPI in NPBT, then washed in NPBT, transferred in a glass-bottomed dish in 100% glycerol and imaged with an Olympus V3000 inverted laser scanning confocal microscope.

#### Zebrafish

Zebrafish embryos were processed as described in (97). Adult zebrafish were decalcified in Morse solution (10% w/v sodium citrate dihydrate and 25% v/v formic acid in DEPC water) prior to embedding for 24h at room temp. Adult and 8 dpf larval zebrafish samples were then cleared with Histosol (National Diagnostics) for 3 x 20min at room temperature, incubated in 1:1 Histosol:Paraffin for 2 x 30min at 60°C, then infiltrated with molten paraffin overnight at 60°C. After additional 4 x 1h paraffin changes, samples were embedded in peel-a-way molds (Sigma), left to set for 24h and then sectioned at 7um on a Leica RM2125 rotary microtome. Sections were mounted on SuperFrost Plus charged glass slides, and *in situ* HCR was then performed as described in (97). Imaging was performed on an Olympus V3000 inverted laser scanning confocal microscope. Images were analyzed using Imaris v.10.0.0 (Oxford Instruments). Due to the high level of autofluorescence from blood and body cavities in whole late embryos, the endoderm was manually segmented using the nuclear staining, and the HCR signal for *FoxQ2c* was then masked within the endoderm to improve visualization (see original image in supplementary fig. 8C).

#### Chicken

Dissected and fixed chicken eyes were rehydrated in PBS and embedded in 4% agarose in PBS. 150µm-thick sections were then obtained using a Leica VT1200S vibratome, and *in situ* HCR on floating sections was performed in a 12-well plate as described in (98). Sections were then mounted on SuperFrost slides with Fluoromount-G (ThermoFisher Scientific 00-4958-02) and imaged using a Zeiss LSM800 inverted laser scanning confocal microscope.

#### Lamprey

For hybridized chain reaction–fluorescence in situ hybridization (HCR–FISH), we adopted the third-generation HCRv3–FISH protocol (95). HCR–FISH probe sets were custom-designed by Molecular Instruments. Following HCR-FISH, embryos and larvae were incubated with SYTOX Green nucleic acid stain (Thermo Fisher Scientific, catalogue no. S7020), followed by brief washes in PBS. Samples were mounted in PBS and then imaged using a Nikon C2 confocal microscope.

#### Skate

Skate S29 embryos were processed for *in situ* HCR as described in (98), with modifications. First, embryos were delipidated by overnight incubation in dichloromethane (DCM). After washes in 100% methanol, the embryonic tail was cut out at the level of the cloaca using razor blades, and samples were rehydrated in PBS and bleached for 30 mins (5% Deionized formamide, 1.5% H2O2, 0.2% SSC in nuclease-free water). Embryos were incubated for 2h with hybridization buffer and then for 3 days with hybridization buffer and 12µM of probes at 37°C. Samples were washed with 5X-SSC + 0.1% Triton X, then amplification buffer, and then left for 3 days at 4°C with 0.15µM of hairpin and 1 µg/mL of DAPI. After incubation, samples were rinsed with 5X-SSC + 0.1% Triton X and then left in TrisHCl 0.5M pH7 overnight. The following day, embryos were embedded in 4% agarose in

TrisHCl 0.5M pH7 and progressively dehydrated in methanol. For clearing, after overnight incubation in 100% methanol, samples were treated with 66% DCM and 33% MeOH for 3h, then with 100% DCM for 1h and then transferred in Dibenzyl ether (DBE) and left overnight. Cleared samples were imaged using an UltraMicroscope II Light-sheet microscope (Miltenyi biotec).

### Transcriptomic data analysis

The following published bulk RNAseq and scRNAseq datasets were used to investigate the expression of *FoxQ2* orthologs across bilaterians:

*Dreissena polymorpha* (Mollusca, Bivalvia): dataset of trochophore larva obtained from (60) available through GEO accession number GSE192624. Normalization (SCTransform), dimensionality reduction and clustering of the raw dataset was carried out using Seurat (99). Marker genes listed in the original publication were used to annotate the clusters.

*Hofstenia miamia* (Xenacoelomorpha): dataset of hatchling juveniles from (61) available in NCBI BioProject database under accession codes PRJNA888438. *FoxQ2* expression was visualized using the online resource generated by the authors https://n2t.net/ark:/84478/d/q6fxc7jj.

*Branchiostoma lanceolatum* (Chordata, Cephalochordata) bulk RNAseq datasets of embryonic and larval development and of adult tissues from (65) were available from GEO with accession number GSE106430. Counts were normalized for library size and composition bias using DESeq2 (100).

*Branchiostoma floridae* (Chordata, Cephalochordata). The single nuclei RNAseq dataset of amphioxus development from (66) was available from CNSA of CNGBdb under accession number CNP0000891. Analysis was performed following the code prepared by the authors and deposited at https://github.com/XingyanLiu/AmphioxusAnalysis.

Seurat objects of processed scRNAseq datasets of specific embryonic stages (B, G0, N0, T1), corresponding to the moment of activation of *FoxQ2a*, *FoxQ2b* and *FoxQ2c* expression in amphioxus, obtained by (69) were downloaded from the Science Data Bank (https://doi.org/10.57760/sciencedb.08801) (101).

*Danio rerio*: bulk RNAseq dataset of embryonic and larval development from (68), stored in the EBI European Nucleotide Archive with accession numbers PRJEB12296, PRJEB7244 and PRJEB12982, was available to explore in Expression Atlas (https://www.ebi.ac.uk/gxa/experiments/E-ERAD-475/Results) (102). Single cell RNAseq of 8 dpf zebrafish larvae from (67) was downloaded from GEO (accession number GSE158142).

All scRNAseq datasets were analyzed and all plots were generated using Seurat v.5.0.3.

### Prediction of regulatory sequences

The genomes of *Branchiostoma lanceolatum* (v.Bl71nemr, GCA_900088365.1), *B. floridae* (Version 2, (103)) and *B. belcheri* (v.Haploidv18h27, GCF_001625305.1), as well as the genomes of *Asymmetron* and *Epigonichthys* available in our labs (Lotharukpong et al., unpublished), were BLAST-searched for the three *FoxQ2* sequences. Then, for *FoxQ2a*- and *FoxQ2b*-type genes, the first 5000bp upstream of each gene’s starting codon sequence were isolated, while for *FoxQ2c*-type genes the upstream 7000bp were selected to account for the large 5’-UTR found in *B. lanceolatum*. For each gene, the five upstream sequences, one for each species, were searched for conserved regions using mVISTA online tool (74) (https://genome.lbl.gov/vista/mvista/submit.shtml), and the regions present in each species were manually annotated using SnapGene Viewer. Moreover, *B. lanceolatum* ATACseq data at four stages of development (8hpf, 15hpf, 36hpf, 60hpf) (65), available at GEO under accession number GSE106428, was mapped on the genome using IGV (104) and the sum of open peaks from all stages was also manually annotated using SnapGene Viewer (supplementary fig. 9). For each gene and each species, the sequences with overlapping sequence conservation for all five species (and, for *B. lanceolatum*, also the sequence contained within the ATACseq peaks) were labelled as conserved non-coding sequences (CNCSs). This resulted in three CNCSs for *FoxQ2a*, one CNCS for *FoxQ2b* and two CNCSs for *FoxQ2c* found in all five amphioxus species. Each CNCS from each species was then analyzed for the identification of putative TFBSs using CiiiDER (75), using JASPAR 2020 core vertebrate matrix of TFBSs (105) and the default deficit threshold of 0.15. This resulted in an extensive list of putative TFBSs, which were then filtered computationally to only include those found in all five species at the same position: the sequence length was normalized between 0 and 1 and TFBS position was used to filter those within the same 10% region for each CNCS (supplementary table 3). This list was then further subset to account for the developmental timing of activation of each *FoxQ2* paralog as well as the cell types in which *FoxQ2* is expressed. We therefore used both the published developmental RNAseq dataset of *B. lanceolatum* from (65) and the developmental scRNAseq of *B. floridae* from (69) (see Transcriptomic data analysis section for details).

- For *FoxQ2a*, in the bulk-RNAseq we averaged the expression of genes at the 32-cells (maternal) and blastula (B, zygotic) stages, while for scRNAseq we selected clusters labelled as “animal pole” at the blastula (B) stage and “epithelial ectoderm” and “ectoderm mix” at the early gastrula (G0) stage, pseudo-bulked using Seurat’s AggregateExpression function, and calculated the average counts for each gene.
- For *FoxQ2b*, in bulk-RNAseq we averaged expression at 11 hpf (G5) and 15 hpf (N0-N2), while for scRNAseq we focused on the N0 scRNAseq dataset, selected clusters labelled as “epithelial ectoderm” and “neural ectoderm”, and calculated the average for each gene of the pseudo-bulked counts.
- For *FoxQ2c* we averaged expression at 36 hpf (T1/14ss) and 50hpf (L0/1gs), and selected clusters within the endoderm where *FoxQ2c* was expressed in the T1 scRNAseq dataset (clusters 5, 8, 67), pseudo-bulked and calculated the average counts for each gene.

For each CNCS, we then subset the list of predicted TFBSs to include only those expressed at the correct developmental time and cell type in which the corresponding gene is expressed. For RNAseq, we considered “significantly expressed” any gene whose expression was higher than the 25th percentile of normalized counts for the whole transcriptome at the stages considered (approximated at 17), while for scRNAseq, to account for the high number of zero counts in different cells, we considered “significantly expressed” genes for which expression was above the 50th percentile (approximated at >250). The final list for each gene is stored in supplementary table 3.

## Supporting information

Gattoni_FoxQ2_SupplementaryFigures

## Acknowledgments

We thank Michael Schwimmer for helping with the analysis of amphioxus adult RNAseq and Colin Shew for providing the object of amphioxus adult scRNAseq dataset. We also thank Jordi Paps for advice and feedback on the phylogenetic analysis, Panagiotis Oikonomou and Nandan L. Nerurkar for providing chicken embryos and Mansi Srivastava for help with the *Hofstenia miamia* dataset.

