## Supplementary material for "Evolutionary dynamics of FoxQ2 transcription factors across metazoans: A tale of three ancient paralogs": Gattoni_FoxQ2_SupplementaryFigures

**Supplementary Figures 1-9**

**Supplementary Tables 1-3 Legends**

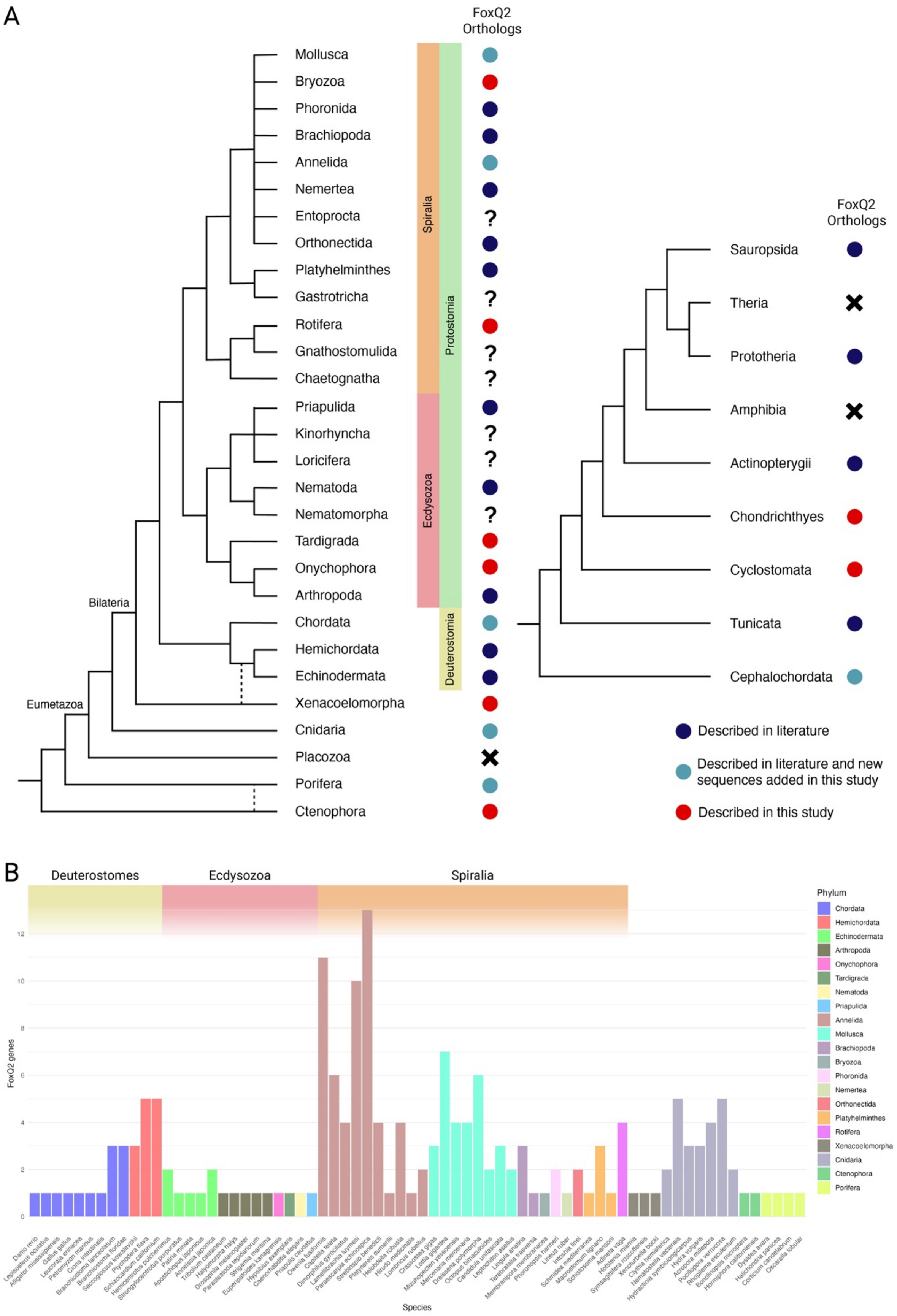
A
FoxQ2 Orthologs
Mollusca
Bryozoa
Phoronida
Brachiopoda
Annelida
Nemertea
Entoprocta
Orthonectida
Platyhelminthes
Gastrotricha
Rotifera
Gnathostomulida
Chaetognatha
Priapulida
Kinorhyncha
Loricifera
Nematoda
Nematomorpha
Tardigrada
Onychophora
Arthropoda
Chordata
Hemichordata
Echinodermata
Xenacoelomorpha
Cnidaria
Placozoa
Porifera
Ctenophora
Spiralia
Ecdysozoa
Protostomia
Deuterostomia
Bilateria
Eumetazoa
FoxQ2 Orthologs
Sauropsida
Theria
Prototheria
Amphibia
Actinopterygii
Chondrichthyes
Cyclostomata
Tunicata
Cephalochordata
Described in literature
Described in literature and new sequences added in this study
Described in this study
B
Deuterostomes
Ecdysozoa
Spiralia
FoxQ2 genes
Species
Phylum
Chordata
Hemichordata
Echinodermata
Arthropoda
Onychophora
Tardigrada
Nematoda
Priapulida
Annelida
Mollusca
Brachiopoda
Bryozoa
Phoronida
Nemertea
Orthonectida
Platyhelminthes
Rotifera
Xenacoelomorpha
Cnidaria
Ctenophora
Porifera

**Supplementary Fig. 1. Conservation of *FoxQ2* genes but variation in copy number across metazoans.**

**A.** Phylogenetic tree showing the relationship between animal phyla, with a focus on chordate evolution. *FoxQ2* orthologs can be identified in most phyla investigated (circles) but appear to be lost in specific lineages (crosses). The color of the circles indicate whether *FoxQ2* sequences for specific phyla were described in the literature (blue), discovered in this work (red), or present in the literature but found for new species in this study (teal). **B.** Barplot indicating the variation in copy number of *FoxQ2* genes (y axis) in each of the 70 species considered in this study (x axis). Bars are colored by phylum.

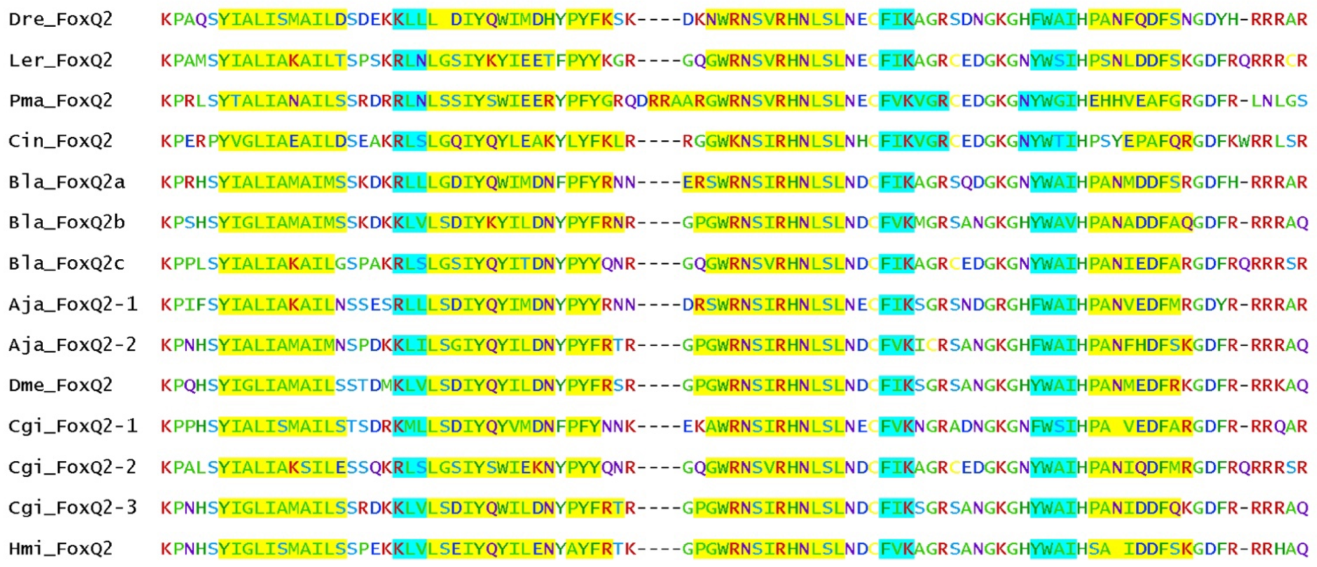


**Supplementary Fig. 2. Broad conservation of predicted secondary structure of the Forkhead domain in representative bilaterian species**.

Amino acids predicted to form alpha-helices are highlighted in yellow, those forming beta-sheets are highlighted in cyan.

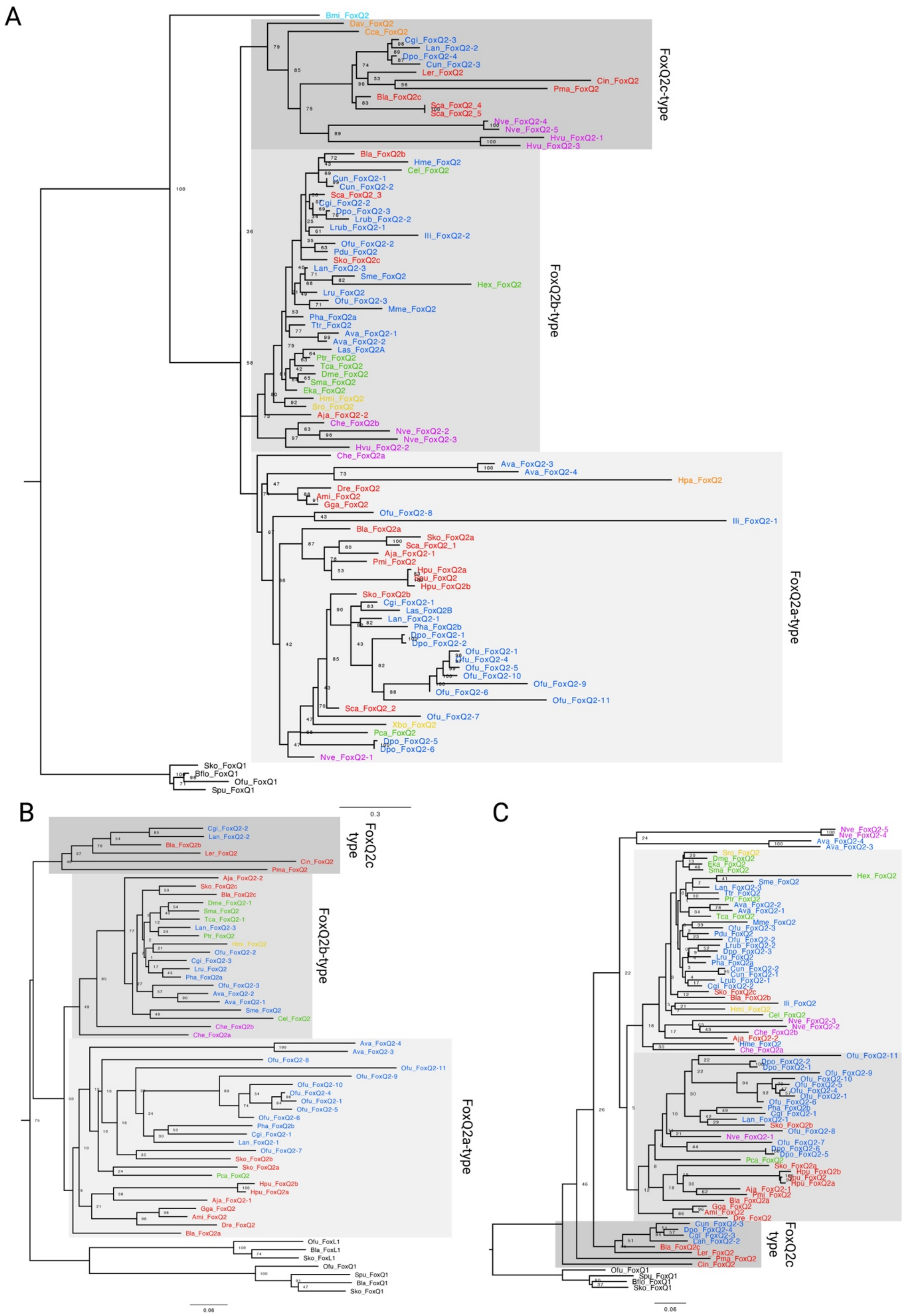
A
FoxQ2c-type
FoxQ2b-type
FoxQ2a-type
0.3
B
FoxQ2c type
FoxQ2b-type
FoxQ2a-type
0.06
C
FoxQ2b-type
FoxQ2a-type
FoxQ2c type
0.06

**Supplementary Fig. 3. Additional phylogenetic trees of FoxQ2 genes across metazoans.**

**A.** Tree obtained with Maximum Likelihood analysis of the Forkhead domain in 47 species from 21 animal phyla. **B-C.** Neighbor joining tree topologies using full *FoxQ2* sequences (**B**) or only the Forkhead domain (**C**). In all trees, sequences are colored based on taxonomy: deuterostomes (red), spiralian (blue) and ecdysozoan (green) protostomes, xenacoelomorphs (yellow), cnidarians (purple), ctenoforans (cyan) or poriferans (orange). Gray-shaded boxes demarcate the three *FoxQ2* types, *FoxQ2a*, *FoxQ2b* and *FoxQ2c*. *FoxQ1* and *FoxP* sequences are used as outgroup.

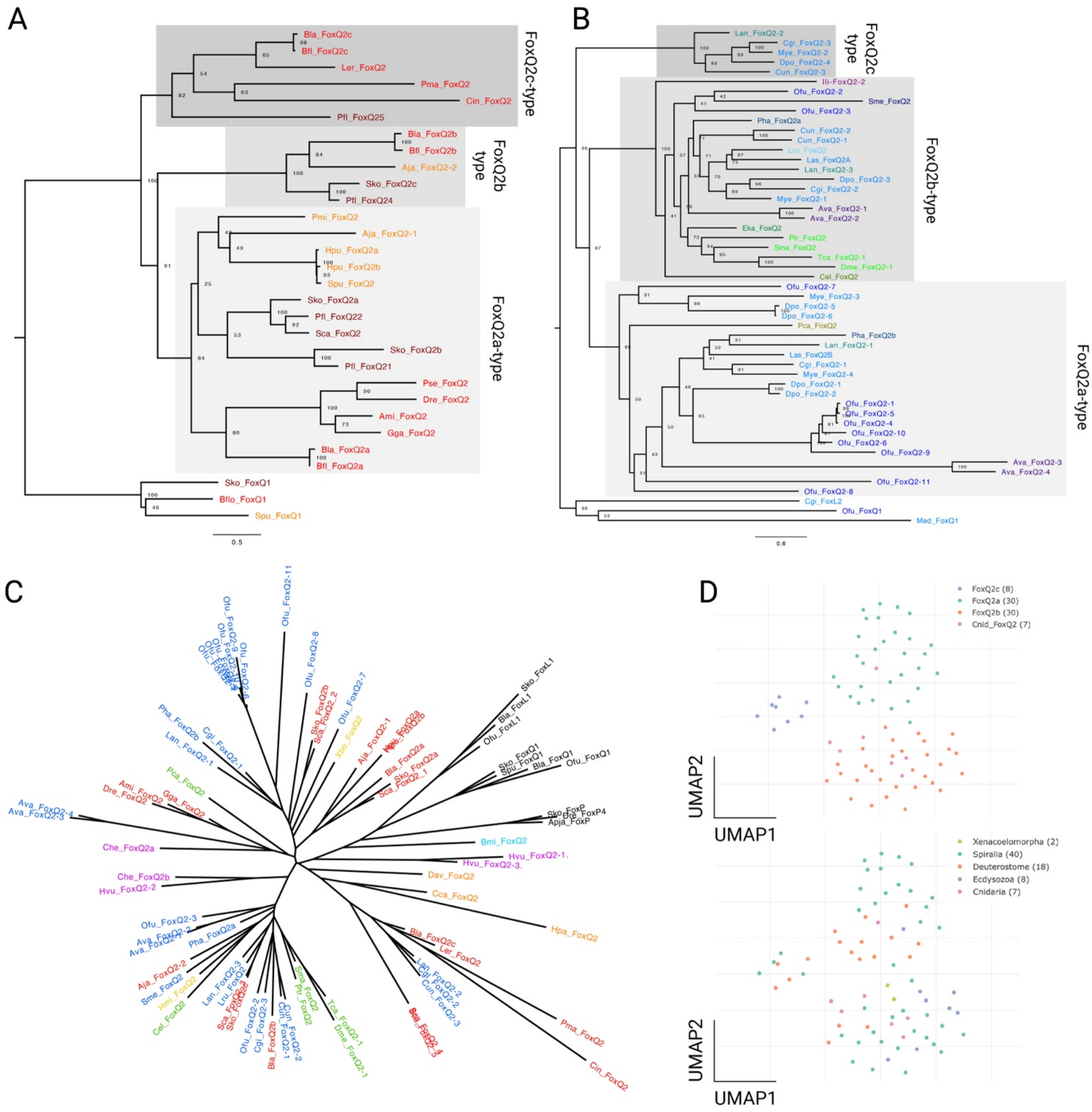


**Supplementary Fig. 4. Additional phylogenetic comparisons of FoxQ2 genes**.

**A-B.** Maximum Likelihood analysis considering only deuterostome (**A**) and protostome (**C**) species recover the same tree topology than the full metazoan phylogenetic tree. Sequences from different phyla are demarcated by different shades of red (A, deuterostomes), blue (B, spiralians) and green (B, ecdysozoa). Gray-shaded boxes demarcate the three *FoxQ2* types, *FoxQ2a*, *FoxQ2b* and *FoxQ2c*. **C.** Unrooted tree resulting from Maximum Likelihood analysis of full *FoxQ2* sequences, showing preservation of *FoxQ2a*, *FoxQ2b* and *FoxQ2c* types separated by dashed lines, and the difficult placement of cnidarian sequences due to high divergence. Sequences are colored based on taxonomy: deuterostomes (red), spiralian (blue) and ecdysozoan (green) protostomes, xenacoelomorphs (yellow), cnidarians (purple). **D.** Result of uniform manifold approximation and projection (UMAP) dimensionality reduction using aligned forkhead domain of metazoan *FoxQ2* genes, colored by *FoxQ2* type (*FoxQ2a*, *FoxQ2b, FoxQ2c* and cnidarian *FoxQ2*) or by taxon, showing the presence of three separated clusters of sequences in accordance with phylogenetic analysis.

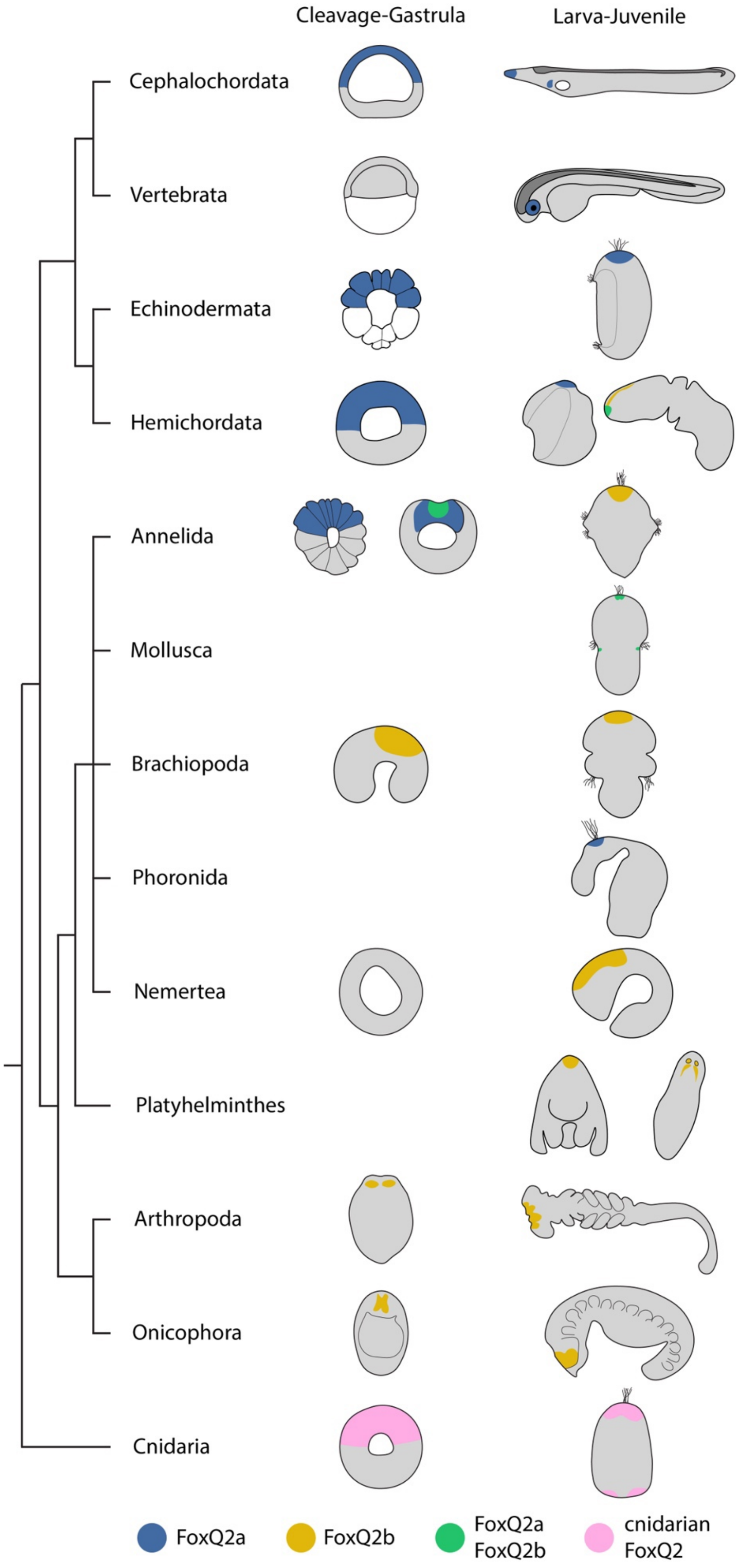

Cleavage-Gastrula
Larva-Juvenile
Cephalochordata
Vertebrata
Echinodermata
Hemichordata
Annelida
Mollusca
Brachiopoda
Phoronida
Nemertea
Platyhelminthes
Arthropoda
Onicophora
Cnidaria
FoxQ2a
FoxQ2b
FoxQ2a
FoxQ2b
cnidarian
FoxQ2

**Supplementary Fig. 5. Summary of *FoxQ2a* and *FoxQ2b* expression during animal development**. Schematic drawings of early and late developmental stages of representative species in which the expression of *FoxQ2a*-type (blue) and *FoxQ2b*-type (yellow) genes has been investigated. Due to the difficult positioning of *FoxQ2a-* and *FoxQ2b*-type genes in cnidarians, the developmental expression domain of *FoxQ2* genes in cnidarians has been colored separately.

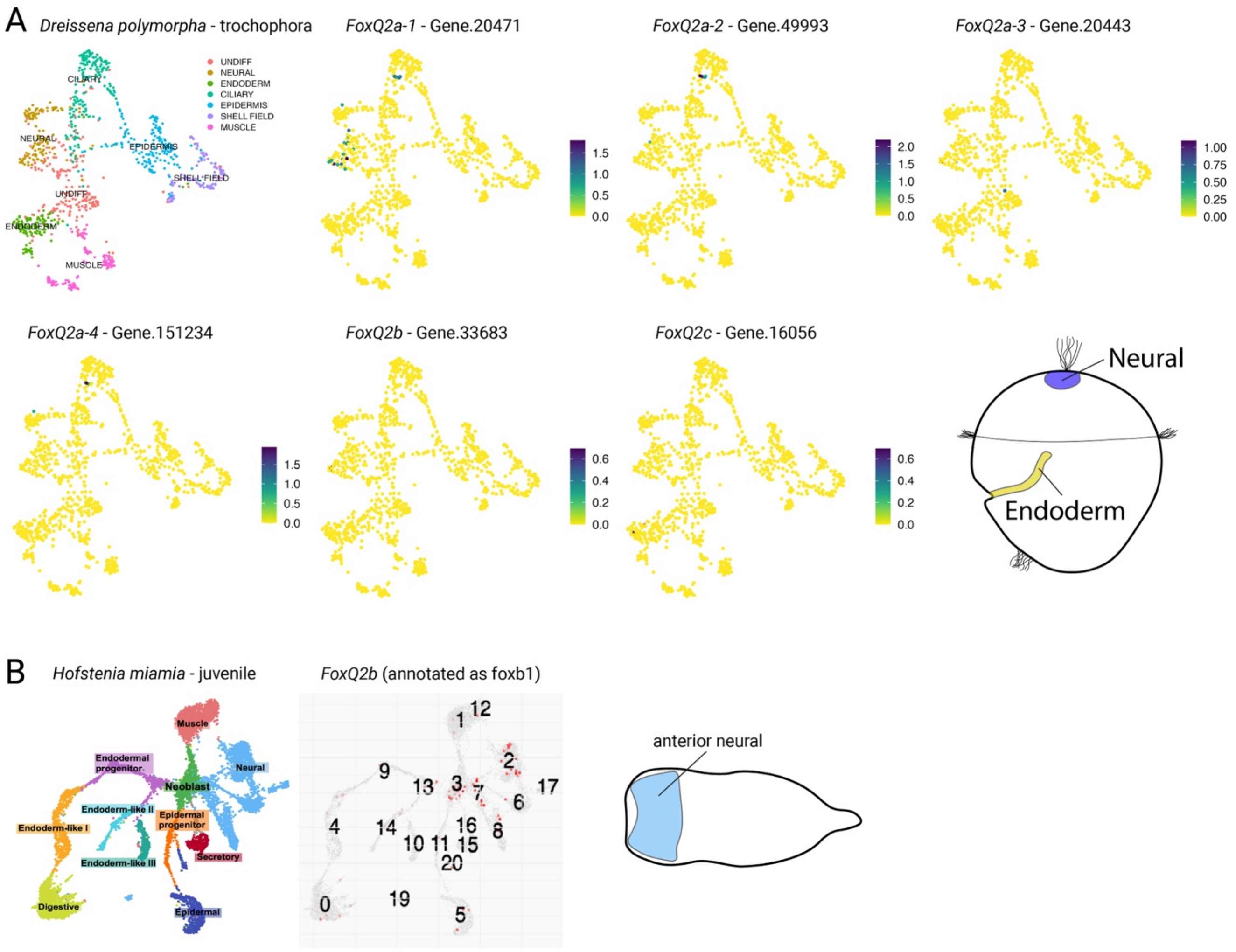


**Supplementary Fig. 6. Analysis of *FoxQ2* expression in publicly available invertebrate single-cell RNAseq datasets.**

**A.** Cluster annotation and expression of *FoxQ2a*, *FoxQ2b*, and *FoxQ2c* orthologs in the early trochophore larva of the bivalve mollusk *Dreissena polymorpha* (data from (60)), showing expression of one *FoxQ2a*-type gene in neurons, sparse expression of two *FoxQ2a* and one *FoxQ2b* gene in ciliated cells, and rare expression of *FoxQ2c* in the endoderm. A schematic representation of the trochophora larva highlights the localization of neural and endodermal cells. **B.** Expression of *FoxQ2b*-type gene in the juvenile stage of the acoel *Hofstenia miamia* (data from(61)), showing positive cells in the neural and neoblast clusters. The *FoxQ2b*-positive neural clusters 2 and 7 have been mapped by the original authors as anterior neural clusters, as represented in the schematic drawing of *H. miamia* juvenile.

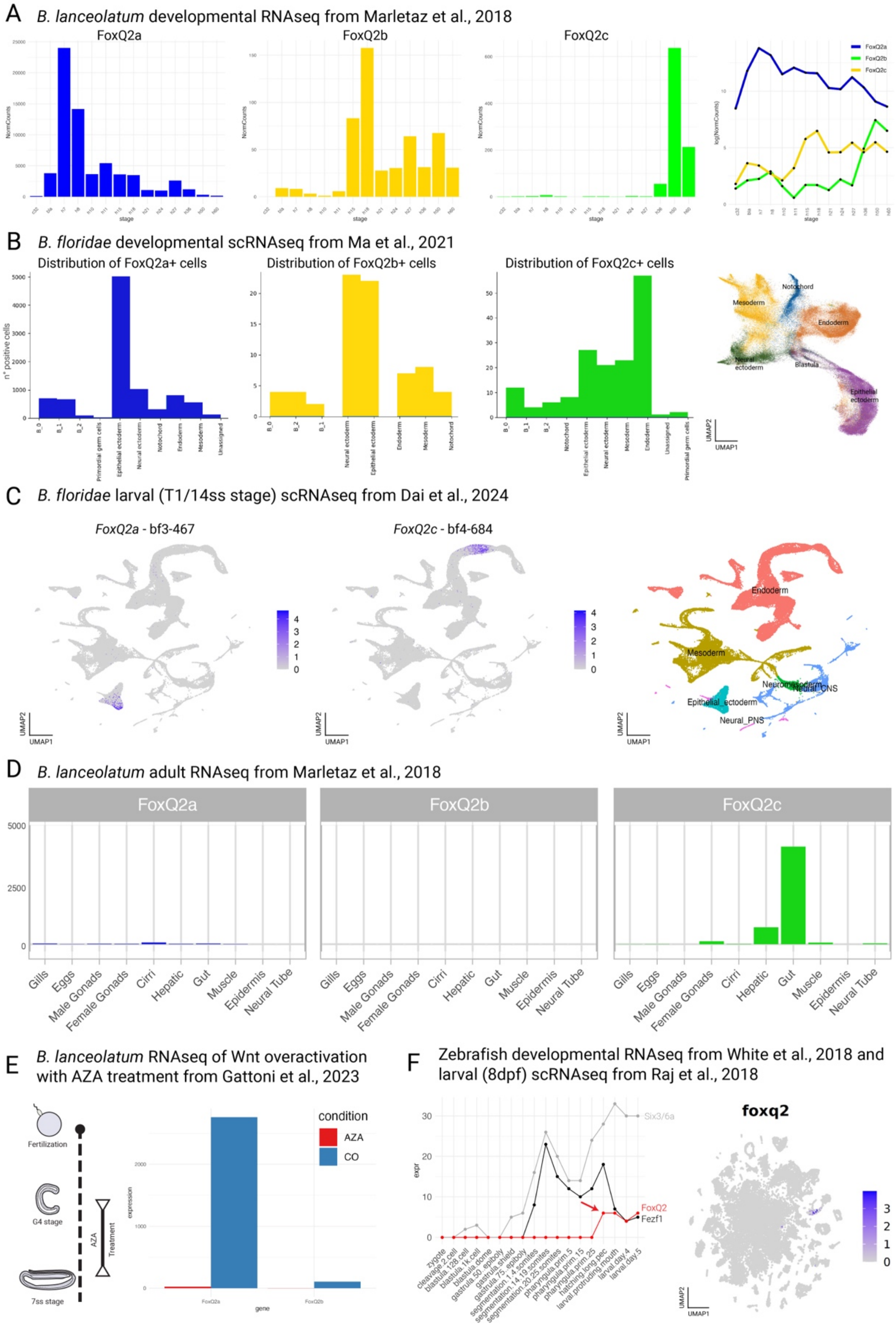

A
B. lanceolatum developmental RNAseq from Marletaz et al., 2018
FoxQ2a
FoxQ2b
FoxQ2c
NormCounts
stage
log(NormCounts)
FoxQ2a
FoxQ2b
FoxQ2c
B
B. floridae developmental scRNAseq from Ma et al., 2021
Distribution of FoxQ2a+ cells
Distribution of FoxQ2b+ cells
Distribution of FoxQ2c+ cells
n° positive cells
B_0
B_1
B_2
Primordial germ cells
Epithelial ectoderm
Neural ectoderm
Notochord
Endoderm
Mesoderm
Unassigned
Notochord
Mesoderm
Endoderm
Neural ectoderm
Blastula
Epithelial ectoderm
UMAP2
UMAP1
C
B. floridae larval (T1/14ss stage) scRNAseq from Dai et al., 2024
FoxQ2a - bf3-467
FoxQ2c - bf4-684
Endoderm
Mesoderm
Neuromesoderm
Neural_CNS
Epithelial_ectoderm
Neural_PNS
UMAP2
UMAP1
D
B. lanceolatum adult RNAseq from Marletaz et al., 2018
FoxQ2a
FoxQ2b
FoxQ2c
5000
2500
0
Gills
Eggs
Male Gonads
Female Gonads
Cirri
Hepatic
Gut
Muscle
Epidermis
Neural Tube
E
B. lanceolatum RNAseq of Wnt overactivation with AZA treatment from Gattoni et al., 2023
Fertilization
G4 stage
7ss stage
AZA
Treatment
expression
condition
AZA
CO
FoxQ2a
FoxQ2b
gene
F
Zebrafish developmental RNAseq from White et al., 2018 and larval (8dpf) scRNAseq from Raj et al., 2018
expr
Six3/6a
FoxQ2
Fezf1
zygote
cleavage 2 cell
blastula 128 cell
blastula 1k cell
blastula dome
gastrula 50 epiboly
gastrula shield
gastrula 75 epiboly
segmentation 1 4 somites
segmentation 14 19 somites
segmentation 20 25 somites
pharyngula prim 5
pharyngula prim 15
pharyngula prim 25
hatching long pec
larval protruding mouth
larval day 4
larval day 5
foxq2
UMAP2
UMAP1

**Supplementary Fig. 7. Additional transcriptomic analyses of amphioxus and zebrafish in support of *in situ* hybridization data.**

**A.** Developmental expression levels of *FoxQ2a* (blue), *FoxQ2b* (yellow) and *FoxQ2c* (green) in *Branchiostoma lanceolatum* from bulk RNAseq (65). Both separated and relative expression levels indicate activation of *FoxQ2a* at the blastula stage, *FoxQ2b* at the early neurula stage and *FoxQ2c* at the early larva stage. **B.** Distribution of *FoxQ2a*, *FoxQ2b* and *FoxQ2c* in single-cell RNA sequencing (scRNAseq) data from amphioxus development (66). The distribution is calculated as the number of positive cells (y axis) in each annotated cluster (x axis), and show expression of *FoxQ2a* and FoxQ2b in neural and non-neural ectoderm and expression of *FoxQ2c* in the endoderm. **C.** This result is further confirmed by the distribution of *FoxQ2a*- and *FoxQ2c*-positive cells in a scRNAseq dataset from *B. floridae* early larvae (69), showing expression of *FoxQ2a* in epidermis and *FoxQ2c* in a specific section of the midgut (endoderm). **D.** Expression level (y axis) of of *FoxQ2a*, *FoxQ2b* and *FoxQ2c* in different tissues of adult amphioxus (65), showing maintenance of high *FoxQ2c* expression in the adult digestive system. **E.** Significant differential expression of *FoxQ2a* and *FoxQ2b* in control (blue) and Azakenpaullone treated (red) embryos, demonstrating loss of expression for both genes following pharmacological overactivation of Wnt signalling. **F.** Bulk RNAseq dataset of zebrafish development (68) shows that *FoxQ2a* starts to be expressed during the beginning of larval stages, much later than the peak of expression of anterior neuroectoderm genes such as *Six3a* and *Fezf1*. scRNAseq data of 8 days old zebrafish larvae (67) shows expression of *FoxQ2* restricted to blue cones and photoreceptor precursors.

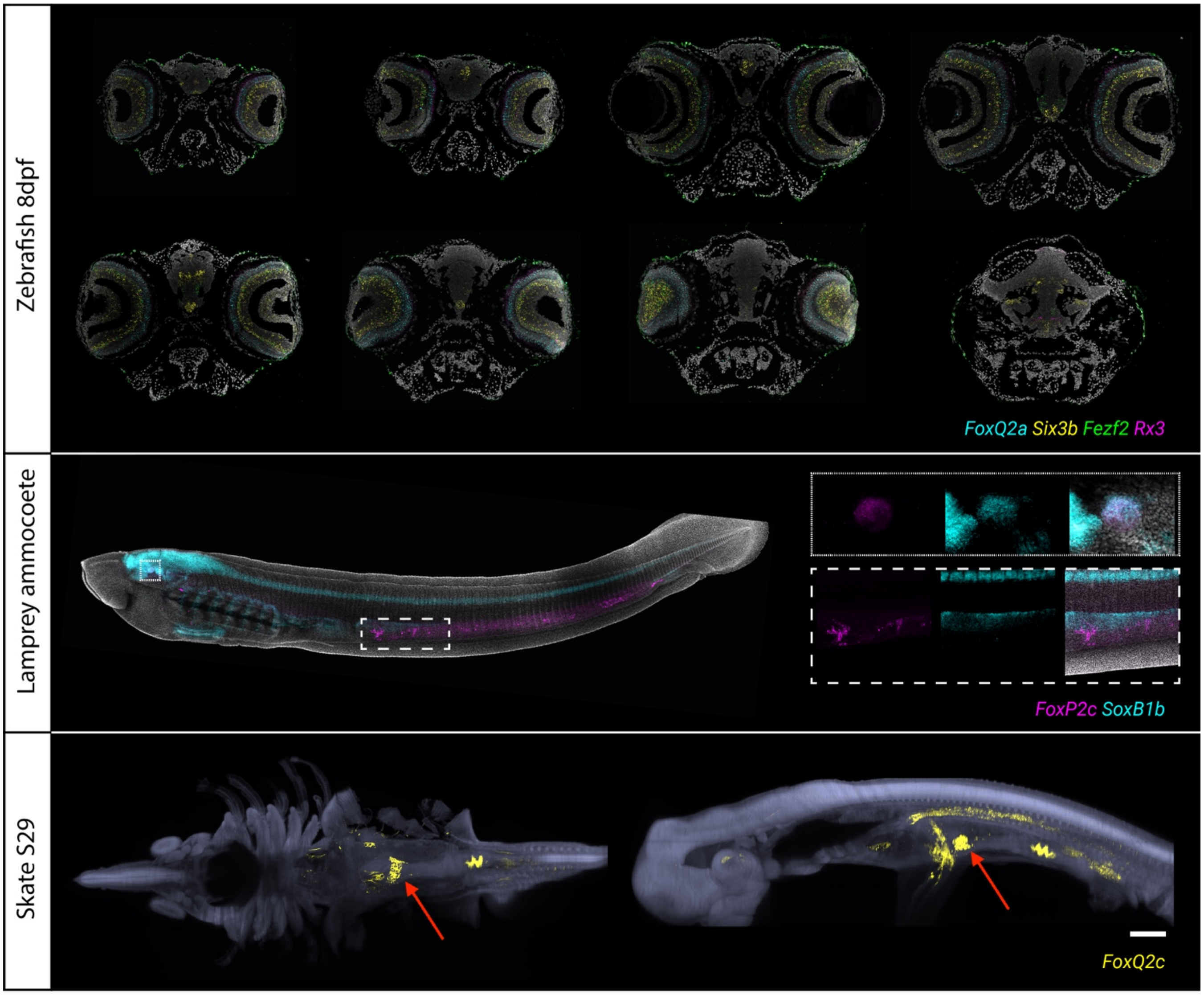


**Supplementary Fig. 8. Additional panels showing expression pattern of vertebrate *FoxQ2* orthologs.**

**A.** Co-detection of *FoxQ2a* with conserved anterior neuroectoderm markers *Six3b*, *Fezf2* and *Rx3* in 8dpf zebrafish larvae. *FoxQ2a* is not expressed in the brain, where all other genes collectively label distinct forebrain cell types, but is expressed in the retina photoreceptor layer. **B.** Whole-mount *in situ* HCR of lamprey ammocoete larva showing expression of *FoxQ2c* and *SoxB1b* in the gut (dashed box) and the eye (dotted box). **C.** Whole-mount S29 skate embryo labelled with *FoxQ2c* before the nuclear masking used in fig. 4, showing aspecific expression in the body cavity. The real expression in the midgut is marked by a red arrow.

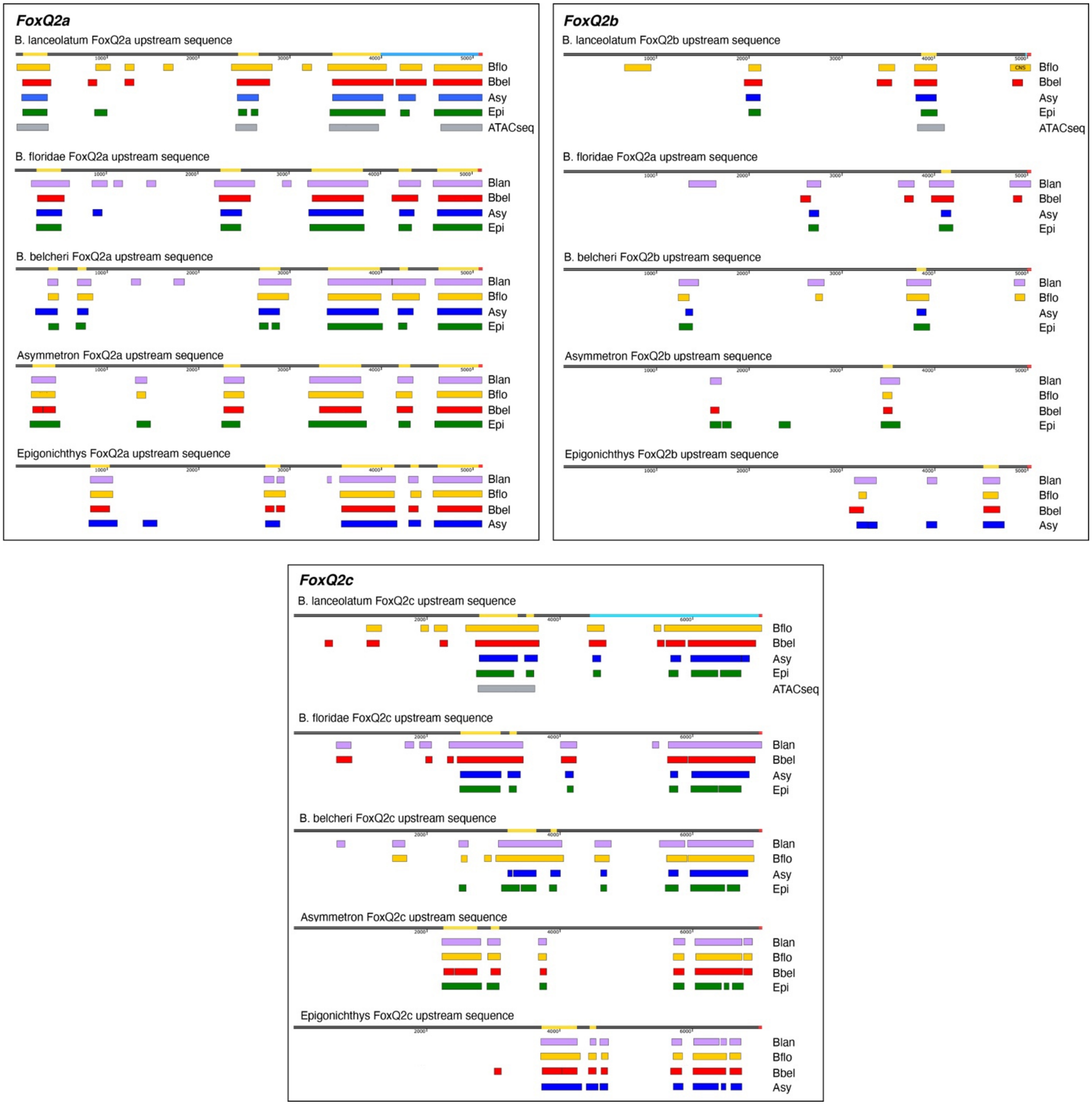


**Supplementary Fig. 9. Conserved Non-Coding Sequences for FoxQ2a, FoxQ2b and FoxQ2c in cephalochordates.**

Conserved nucleotide sequences upstream of the three *FoxQ2* genes (*FoxQ2a*, *Fox2b*, *FoxQ2c*) in five cephalochordate species identified using mVISTA. Each *FoxQ2* upstream sequence for each species was compared to all other four species. For each sequence (represented as a black double line), conserved non-coding sequences (CNCSs) are highlighted in yellow, the starting codon is in red and the 5'UTR region is in light blue. For each species, the conserved regions are indicated with violet (*Branchiostoma lanceolatum*), yellow (*B. floridae*), red (*B. belcheri*), blue (*Asymmetron*) and green (*Epigonichthys*) bars. CNCSs are defined as the regions that show conservation across all five species. For *B. lanceolatum*, the open chromatin regions identified by ATACseq are marked by a grey bar, showing that they correspond to the CNCS regions.

**Supplementary table S1. Complete list of FoxQ2 genes in 70 metazoan species, with division into a, b and c types and new nomenclature**

**Supplementary table S2. Macro-synteny analysis results for 29 metazoan species and two non-metazoan Opisthokonta outgroups.**

**Supplementary table S3. Analysis of Conserved Non-Coding Sequences (CNCSs) in cephalochordates. The table contains:**

- **Sequences of each CNCS in *B. lanceolatum***
- **List of conserved amphioxus transcription factors that have identified transcription factor binding sites (TFBSs) in each CNCS and that are expressed together with each FoxQ2 gene**
- **List of transcription factor families identified in the study**
